# LIET Model: Capturing the kinetics of RNA polymerase from loading to termination

**DOI:** 10.1101/2024.10.03.616401

**Authors:** Jacob T. Stanley, Georgia E.F. Barone, Hope A. Townsend, Rutendo F. Sigauke, Mary A. Allen, Robin D. Dowell

**Affiliations:** BioFrontiers Institute, University of Colorado Boulder, Boulder, 80303, CO, USA; Molecular, Cellular and Developmental Biology, University of Colorado Boulder, Boulder, 80309, CO, USA

**Keywords:** Transcription regulation, Modeling, Nascent-sequencing, Bioinformatics

## Abstract

Transcription by RNA polymerases is an exquisitely regulated step of the central dogma. Transcription is the primary determinant of cell-state, and most cellular perturbations impact transcription by altering polymerase activity. Thus, detecting changes in polymerase activity yields insight into most cellular processes. Nascent run-on sequencing provides a direct readout of polymerase activity, but no tools exist to model this activity at genes. We focus on RNA polymerase II—responsible for transcribing protein-coding genes. We present the first model to capture the complete process of gene transcription. For individual genes, this model parameterizes each distinct stage of transcription—*Loading, Initiation, Elongation*, and *Termination*, hence LIET—in a biologically interpretable Bayesian mixture, which is applied to nascent run-on data. Our improved modeling of *Loading* /*Initiation* demonstrates these are characteristically different between sense and antisense strands. Applying LIET to 24 human cell-types, our analysis indicates the position of dissociation (the last step of *Termination*) appears to be highly consistent, indicative of a highly regulated process. Furthermore, applying LIET to perturbation experiments, we demonstrate its ability to detect specific changes in pausing (5***′***end), strand-bias, and dissociation location (3***′***end)—opening the door to differential assessment of transcription at individual stages of individual genes.

## 1 Introduction

RNA polymerases (RNAPs) are the cellular machinery directly responsible for the production of essentially all RNA molecules from DNA within the cell—a process referred to as transcription. Transcription is a key driver of development and a cell’s response to the environment. In order to serve this function, transcription must be intricately regulated. RNA polymerase II (RNAP2) transcribes the largest fraction of the genome, including protein-coding genes, long noncoding RNAs, and enhancer-associated RNAs. The process of transcription by RNAP2 follows a well characterized cycle that includes four sequential phases: loading, initiation, elongation, and termination[1, 2, 3, 4].

Transcription is regulated through mechanisms that impact how RNAP2 is distributed across the genome. Understanding these mechanisms requires detecting changes, sometimes subtle, in the activity of RNAP2 between conditions. Nascent run-on sequencing assays—precision run-on sequencing (PRO-seq[5]) and global run-on sequencing (GRO-seq[6])—produce a direct measure of the distribution of active RNAP2 across the genome, by enriching for the newly synthesized RNA molecule still attached to the polymerase[7]. When generated from a statistically representative population of cells and to sufficient depth, these assays are capable of capturing and quantifying the impacts of regulatory mechanisms on RNAP2 activity, regardless of whether the impacts are extensive or subtle, genome-wide or gene-specific[7, 8]. Consequently, powerful analytical tools have been developed to capture and quantify the patterns of reads present within nascent run-on data.

Each newly developed model for analyzing RNAP2 activity has uncovered new regulatory mechanisms. The earliest analysis efforts focused only on locating regions of active transcription[9, 10, 11, 12], uncovering extensive nascent transcription genome-wide. These earliest studies uncovered long stretches of transcription downstream of the cleavage and polyadenlyation (PAS) site at all genes[9], resulting from continued RNAP2 activity that spatially separates the location of RNA cleavage from RNAP2 termination and dissociation, further along the DNA. However, these early approaches did not leverage the unique profile of reads inherent to each stage of RNAP2 activity and observable in nascent run-on assays. Subsequently, methods were developed to capture the unique bidirectional peak signal observed (loading and initiation[12, 13, 14, 15, 16]), uncovering tens of thousands of transcribed regulatory elements (TREs) that could be leveraged to infer transcription factor activity[15, 17, 18, 19, 20, 21]. Others have focused more on the transition from initiation to elongation (pausing) [2] or the rate of transcription through the gene (elongation)[2, 10, 22]. Over time, the emphasis shifted from pattern detection efforts to richer modeling of the unique activity of RNAP2[14, 22, 23]. However, despite the intriguing stretches of transcription after the PAS, the termination stage of transcription has been largely ignored in RNAP2 modeling.

Regardless of which modeling approach is employed, the regions of RNAP2 activity identified are subsequently quantified. The most common approach to quantification is to count nascent run-on reads over the region, like a gene body, and then use these counts to compare samples/conditions using differential assessment tools like DESeq2[24]. Counts based approaches identify statistically significant changes on a per-gene basis but are insensitive to the complex distribution of active RNAP2 within the region. Consequently, the other common quantification approach uses meta-gene profiles. Meta-genes are generated by aligning the read profiles from many genes by a given coordinate, typically their annotated transcription start site (TSS). In the meta-gene approach, profiles are then compared either between gene sets or between samples/conditions. Meta-genes enable detailed comparison of the RNAP2 distribution of regions between samples/conditions, but can be influenced by the point of reference used to align the genes. Furthermore, gene-specific effects are averaged away, making it unclear to which genes any observed differences can be attributed. For example, Integrator is a protein complex that plays an important role in regulating RNAP2 pausing/elongation and is controlled through interactions with various transcription factors. A meta-gene analysis of a knock-down of Integrator’s endonuclease subunit showed inhibition of RNAP2 release from 5′ pausing[25]. Due to Integrator’s ubiquity at protein-coding genes, it is assumed that this behavior is universal, but it remains possible that the strength of this behavior varies dramatically between genes and some genes may escape this inhibition altogether. What is needed is a rigorous method of comparing the distribution of reads (i.e. the shape of the data) across samples on a per-gene basis.

To address this challenge, we developed a computational model, rooted in the molecular activity of RNAP2 during transcription, that could be applied to individual genes within nascent run-on sequencing data. Our model builds on our previous modeling framework[23], but includes an explicit model of *Termination*. Hence the model effectively captures all stages of RNAP2: *Loading, Initiation, Elongation* and *Termination* on both strands, and is thus called LIET. Furthermore, LIET is flexible, efficient, and capable of leveraging known prior information (when available) yet powerful enough to be driven by the data when it conflicts with our prior expectations. This flexibility also includes the ability to independently model the sense and antisense-strands of the 5′ transcription profile. Importantly, the LIET framework enables gene-specific assessment of changes in RNAP2 activity, which manifest within the data as changes in the shape of read distributions. Here we describe the design and technical details of LIET and assess its ability to detect subtle changes to RNAP2 profiles at both the 5′end (e.g. changes in RNAP2 pausing) and the 3′end (e.g. extension in downstream run-on transcription). Ultimately, the LIET model proves to be an unparalleled tool for analyzing the complexities of nascent run-on sequencing data, opening new avenues for understanding the regulation of transcription.

## 2 METHODS AND MATERIALS

### 2.1 LIET Model: mathematical description

The LIET model is a generative, probabilistic mixture model that captures the four stages of transcription on the sense-strand of the gene, and independently the *Loading* and *Initiation* stages on the antisense-strand, from nascent run-on sequencing data. For *Loading, Initiation*, and *Termination*, the LIET model utilizes well-established probability distributions, with all components defined as functions of the genomic coordinate *z* (i.e. *z* is a relative coordinate, measured relative to the gene TSS, and thus defined on the integers—*z* ∈ ℤ).

The mathematical representations used for *Loading* and *Initiation* are the same as previous work[23]. Briefly, the *Loading* position is treated as a random variable *L*, modeled as a normal distribution with location *μ*_*L*_ and uncertainty *σ*_*L*_, and the *Initiation* distance (a.k.a. “entry length”[4]) is a random variable *I*, modeled as an exponential distribution with characteristic length *τ*_*I*_ (see Eq. 1 and the graphical representation in Fig. 1A).

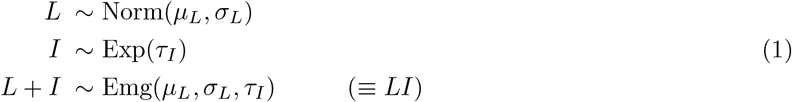

The *Loading* stage is not a directly observable quantity as RNA is not produced from RNAP2 during loading. Because the *Loading* stage must be immediately followed by the *Initiation* stage, it is useful to consider the linear combination of the two into a single random variable within the model, *L* + *I* ≡ *LI*. The convolution of these two produce an exponentially modified Gaussian[23, 26] (a.k.a. ‘Emg’ in Eq. 1, blue/red components—positive/negative strand—in Fig. 1C), whose probability density function (*pdf*) is given by Eq. 4. This combined variable *LI* is a component of the mixture model.

**Fig. 1:**
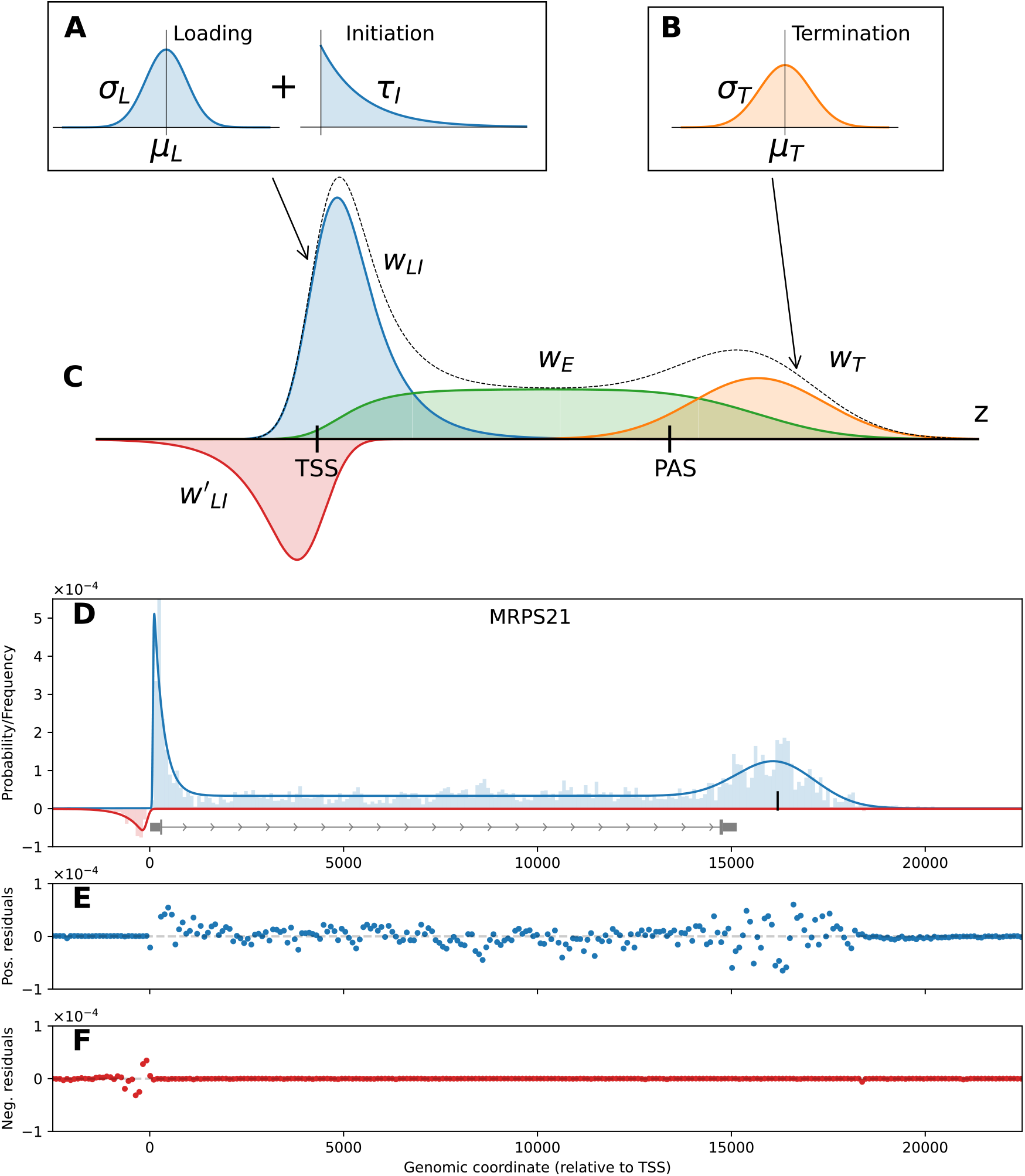
The LIET model. The distributions used to define the processes (A) *Loading, Initiation* (blue), and (B) *Termination* (orange) respectively. (C) The full LIET model with each of the separate components shown, along with the combined distribution (black dotted line). The *Elongation* region (green) is defined based on the two components that flank it (*Loading*+*Initiation* in blue; *Termination* in orange), described fully in Section 2.1. The full model also includes an upstream antisense *Loading* +*Initiation* region, in red. (D) An example of the full model fit to MRPS21 from HCT116 data. The (E) positive and (F) negative strand residuals for MRPS21 showing that the data is well captured at both the 5’ and 3’end of the profile.

Transcription *Loading* +*Initiation* is predominantly bidirectional. To capture this phenomena, our previous model defined a single exponentially modified Gaussian and used an indicator function to specify the strand[23]. However, this tied together all parameters describing the loading and initiation stages of RNAP2. However, we have observed that the shape of the sense and antisense components of bidirectional profiles at the 5′end of genes (*Loading* +*Initiation*—blue and red components in Fig. 1C, respectively) rarely appear to be equivalent in nascent run-on sequencing assays (see example gene profiles in Fig. 2A). Therefore, we sought a more flexible modeling framework to capture potentially distinct shapes on each strand. To this end, the LIET model represents *LI* on each strand independently with *p*_*LI*_ (·) (Eq. 4)—sense parameters: (*μ*_*L*_, *σ*_*L*_, *τ*_*I*_); antisense parameters: 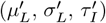. In our implementation, the two strands can then be explicitly tied together to mimic our previous work, associated through shared priors, or treated fully independently.

**Fig. 2:**
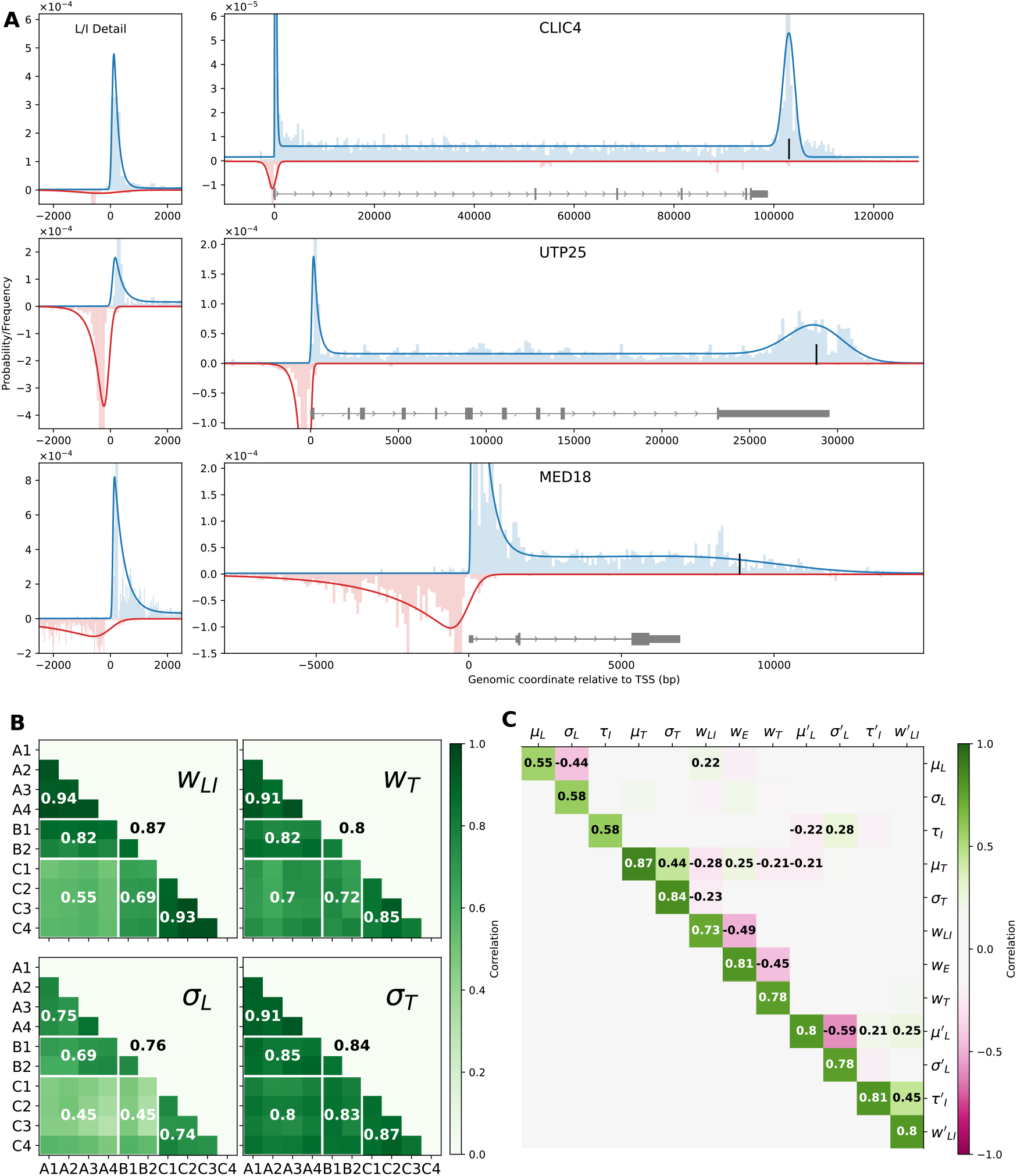
LIET reproducibly fits a variety of gene profiles. (A) Three genes (CLIC4, UTP25, and MED18) highlight the variety of gene lengths and data profiles. (B) Correlation between model on ten control replicates of HCT116 data from three papers [28, 29, 30], designated A/B/C with 4/2/4 replicates, respectively. The numbers in each heatmap are the average correlations for whole-publication comparisons (A:A, A:B, B:C, C:C, etc—delineated by white lines). (C) Median cross-parameter correlation coefficients, with only those whose magnitude is *>* 0.2 are labeled. This correlation matrix is reproduced in greater detail in Supp. Fig. 9.

Once *Initiation* is complete, RNAP2 transitions into the *Elongation* stage of transcription. As we assume RNAP2 stages are ordered both temporally and spatially, we make two key observations on which the mathematical derivation of the *Elongation* component of the LIET Model is based:

- **Observation 1:** The fraction of active RNAP2 that may be in the *Elongation* stage, at a given genomic location is **proportional to** the fraction of the population that have already undergone *Loading* +*Initiation* **upstream** of this location.
- **Observation 2:** The fraction of active RNAP2 that may be in the *Elongation* stage, at a given genomic location is **proportional to** the fraction of the population that will undergo *Termination* **downstream** of this location.

Thus our definition of the *Elongation* stage depends both on how we model *Loading* +*Initiation* and the *Termination* stages.

In *Termination* we seek to capture the furthest 3′-extent to which RNAP2 transcribes beyond the end of the gene, as well as the distribution of signal at that end of the transcriptional profile. In effect, since this is where RNAP2 stops transcribing, it is also the location of dissociation of RNAP2 from the DNA template. Hence, we refer to this process as both transcription termination and RNAP2 dissociation. It is important to note this is distinct from the cleavage and polyadenylation of the mRNA, which some label as “termination” of the transcript—a signal observable in RNA-seq data but not in nascent run-on sequencing. We assume that dissociation occurs downstream of a fixed point, typically a point upstream of the cleavage and polyadenlyation site (PAS). Thus, we treat the termination process (Fig. 1B) as a random variable *T* which we assume, similar to the *Loading* stage, is a symmetric, peaked distribution centered on the genomic dissociation location. This distribution is selected to enable capturing of the apparent 3′end peak commonly observed in nascent run-on sequencing data at protein-coding genes, downstream of the cleavage site[27] (see examples of this in Fig. 1D and 2A). Thus we model *T* as a Gaussian distribution downstream of the PAS with mean *μ*_*T*_ and standard deviation *σ*_*T*_ —*pdf* given in Eq. 6. Notably, the prominent peak in the signal downstream of the PAS is commonly thought to result from RNAP2 slowing prior to dissociation, so we will refer to *μ*_*T*_ as the position of dissociation and the variance *σ*_*T*_ as the fidelity or spread of the dissociation process. It should also be noted, since *T* is a component of the entire mixture model, the amplitude of *p*_*T*_ (·) is an adjustable parameter (*w*_*T*_ in Fig. 1C), which allows LIET to capture dissociation peaks of any prominence (see top and bottom examples in Fig. 2A).

Now that we have described the model components at the two ends of the profile, we return to *Elongation* (green in Fig. 1C), whose mathematical formulation is derived from that of the *LI* and *T* variables. We define the *Elongation* component as a random variable *E*. To conform to the constraints (Observations 1 and 2), a wholly novel probability distribution needed to be derived. Mathematically, Observation 1 implies that the probability distribution of *E* must be proportional to the cumulative distribution function (*cdf*) of *LI* distribution: P(*LI* > *z*) = ∫^*z*^ *p*_*LI*_(ζ) dζ, where *p*_*LI*_ (·) is given by Eq. 4. Observation 2 implies the distribution of *E* is also proportional to the survival function of the *T* distribution: 1 *−* P(*T > z*) = 1 *−* ∫^*z*^ *p*_*T*_(ζ) dζ, where *p*_*T*_ (·) is given by Eq. 6. Combining these two constraints results in *p*_*E*_ (·) in Eq. 2:

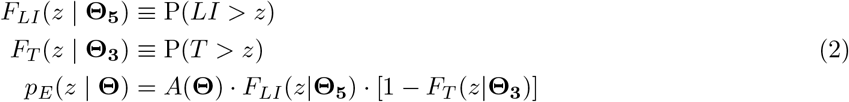

where *F*_*LI*_ (·) and *F*_*T*_ (·) are the *cdf* of the subscript variables, **Θ**_**5**_ and **Θ**_**3**_ are the respective 5′/3′end parameters, and *A*(**Θ**) is a yet undetermined normalization constant that depends on all (5′ and 3′) model parameters (**Θ** = [*μ*_*L*_, *σ*_*L*_, *τ*_*I*_, *μ*_*T*_, *σ*_*T*_ ]).

The key to establishing *p*_*E*_(·) as a proper, functional probability distribution was finding a solution to the normalization constant *A*(**Θ**) in Eq. 2. For this, it is necessary to solve the integral in Eq. 3

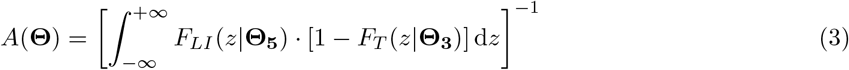

(defined on *z ∈* ℝ). Unfortunately, there is no known closed-form solution (or even reduced-form solution) for integrals of this type in standard integral tables (see ref. [31]). Since genomic position is defined on the integers (ℤ), we initially approached this problem by performing numeric integration of the normalization constant over a finite range of ℤ anytime *p*_*T*_ (·) gets evaluated (numeric approximation method described in Supplemental Sec. 1.1 and Supp. Fig. 1). However, this approach required significant computation, making it infeasible in high-throughput use. Conversely, we were able to derive a partial analytic solution to the integral in Eq. 3 in which the only non-solvable terms were two evaluations of the standard normal *cdf* (Φ(·)). This “partial analytic” solution proved to be approximately 1000x faster on average (see Supp. Sec. 1.4) than the numeric integration method. The analytic method was bench-marked and tested for precision against the numeric method, which we show to be equivalent to high-precision (*<* 10^*−*5^, see Supp. Fig. 2 and 3). For a complete derivation of the partial analytic solution to the normalization constant, see Supp. Sec. 1.2. The explicit mathematical form of *p*_*E*_() is stated in Eq. 5 and the solution to the normalization constant *A*(**Θ**) is Supp. Eq. S.20. Our software implementation of the model uses the analytically normalized form for the *Elongation* component.

As all sequencing data contains noise, we also include an explicit *Background* component, which we treat as a random variable *B* that is modeled as a uniform distribution (independently, on each strand) over the width of the fitting window (*p*_*B*_(·) in Eq. 7). Importantly, the *Background* is not considered part of productive transcription for the gene but rather represents the low-level random read noise, typically ubiquitous throughout the genome in nascent run-on data. Because the position of *μ*_*T*_ is not known *a priori*, the inclusion of the *Background* component also limits the possibility of random read-mapping leading to bias in the *Termination* component. Additionally, including the background component was found to improve fit convergence and quality (not shown). Generally, the background appears to be around 1–5% of the reads when fitting most genes with significant signal.

Finally, the complete model consists of a weighted mixture of these components—*LI, E, T*, and *B* on the sense strand and *LI′* and *B′* on the antisense-strand. The weight vectors are generated from four- and two-dimensional Dirichlet distributions for the sense and antisense mixtures, respectively (sense weights: **w** = [*w*_*LI*_, *w*_*E*_, *w*_*T*_, *w*_*B*_] and antisense weights: 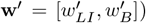. Conceptually, the weights also have the advantage of capturing the relative levels of each component—in other words, multiplying the weights by the total reads within the fitting window produces the number of reads that belong to the respective transcriptional stage, akin to counting reads over exons from RNA-seq data (see discussion in Supp. Sec. 3). The full sense and antisense LIET model likelihood functions are given by Eq. 8 and 9, respectively. Example fits of the full model can be seen in Fig. 1D and 2A.

### 2.2 Model inference and prior selection

Next we sought to provide an easy-to-use software implementation of the LIET model and demonstrate its effectiveness on a number of nascent run-on sequencing datasets. Our software implementation uses a Bayesian inference framework in which the parameters’ priors are informed by gene annotation.

Specifically, for a single gene fit, given a set of observations **X** = *{x*_1_, *x*_2_, …, *x*_*n*_*}* (i.e. a list of relative genomic coordinates of sense-strand reads, *x*_*i*_ *∈* ℤ), the Bayesian inference process amounts to approximating the posterior distribution *p*(**Θ, w** | **X**) from Eq. 10,

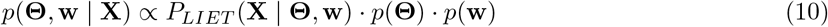

where *P*_*LIET*_ (·) is the sense-strand likelihood function (Eq. 8), *p*(**Θ**) is the prior distribution for the sense-strand model parameters, and *p*(**w**) is the prior distribution for the component weights. An equivalent inference problem exists for the antisense-strand, with data **X**′ and posterior *p*(**Θ**′, **w**′ | **X**′).

The use of priors is key to the success of our model implementation and is another feature that differentiates it from previous nascent run-on sequencing analysis tools. The principle benefit to using a Bayesian approach to model fitting is that the priors focus the parameter search on the most relevant portion of the parameter space. In other words, the priors guide the parameter values to the ranges that are biologically relevant. This has the effect of improving fit convergence and reducing the chance that fits gets stuck in local minima of the parameter landscape. In general, poor fit convergence is exacerbated by sparse/low-coverage data and most nascent run-on sequencing datasets are of low-coverage[17]. Therefore, the use of priors improves the success of fitting nascent run-on data, in particular. In our software implementation, we utilize variational inference (VI) (specifically, Automatic Differentiation Variational Inference[32, 33]) for model fitting. We selected a VI technique for speed of fitting, as our goal is to be able to apply the model to a large number of genes and data sets. One limitation of VI methods is their underestimate of variance of the posterior density[32], which is true in our case as well—see Supp. Figs. 4 and 5. However, the model is separate from the optimization method and can also be used with other VI or MCMC methods.

Model components:

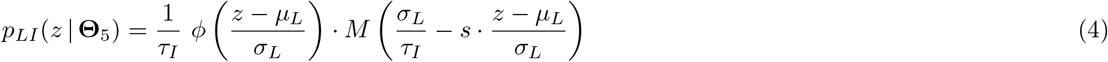

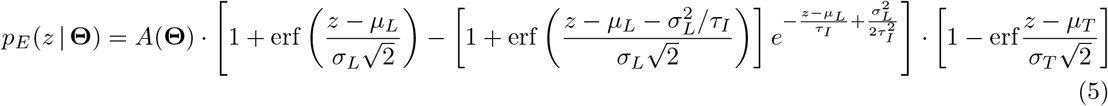

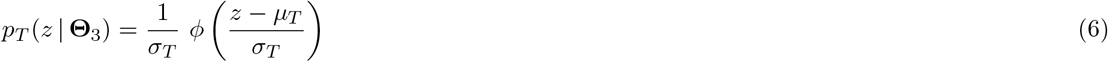

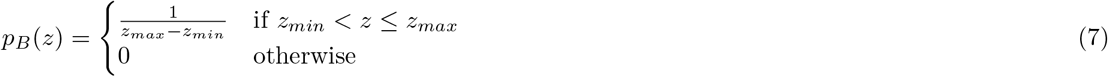

Full model likelihood (sense-strand):

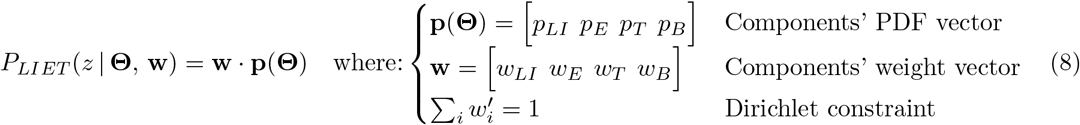

(antisense-strand):

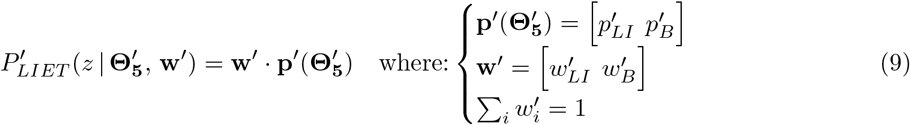

Complete mathematical details for the LIET model. Note: **Θ**_**5**_ = *μ*_*L*_ *σ*_*L*_ *τ* _*I*_, **Θ** _**3**_ = *μ*_*T*_ *σ*_*T*_, and **Θ** = **Θ** _**5**_ **Θ** _**3**_ (antisense: 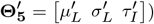). All model components are defined on (relative) genomic coordinates (i.e. *z ∈* ℤ). The normalization constant *A*(**Θ**) for the *Elongation* component is defined in Supp. Eq. S.20. The *pdf* for components *Loading* +*Initiation* (blue/red), *Elongation* (green), and *Termination* (yellow) in Fig. 1C are given by equations 4, 5, and 6, respectively. The Mills ratio: *M* (*·*) = (1 *−* Φ(*·*))*/ϕ*(*·*), where *ϕ*(*·*) is the standard normal distribution and Φ(*·*) is its cumulative distribution function. The strand indicator *s ∈ {*+1, *−*1*}* denotes the positive or negative strand.

For our purposes, the practical goal of fitting our model to real data is to obtain best estimates for each model parameter 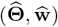, for each gene we fit, so we do not need the detailed shape of the posterior distributions (further justification for using the simpler, quicker VI methods). For a single fit, the parameters’ maximum likelihood estimates (MLEs) are computed as the expectation of the posterior distribution (Eq. 11, and equivalently for **ŵ**). These MLE values are what are reported in the results files.

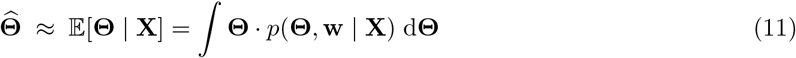

Another common issue in modeling that can impact fitting and the interpretation of results is the presence of confounded parameters. For the LIET model, we observe that there is little to no correlation between the posteriors (Supp. Fig. 9), so the parameters can be treated as essentially independent and thus the posteriors are not likely to be complex/multi-modal. Since the parameters are effectively independent (i.e. not confounded) the prior distribution can be computed by Eq. 12.

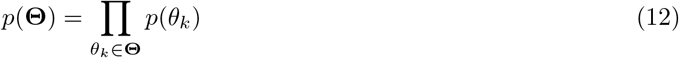

The model parameters consist of three conceptual categories: location parameters 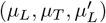, shape parameters 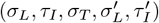, and weight parameters (**w, w′**)—see model diagrams in Fig. 1A–C. The priors for the location parameters leverage reference points to guide their optimization. Specifically, for the sense-strand, the user must provide both 5′ and 3′ end reference points for each gene (*z*_5_, *z*_3_), and the distribution of the priors for the loading and termination location parameters are then built around these points. Conceptually, the *z*_5_ coordinate (which anchors the search for *μ*_*L*_, *μ*_*L*_′) is expected to be in close proximity to the gene’s transcription start site (TSS), whereas the *z*_3_ coordinate (which anchors the search for *μ*_*T*_) is assumed to be proximal to the annotated PAS or some other point guaranteed to be upstream of the dissociation position (*μ*_*T*_).

Hence, in this work, we assume the TSS location will be the 5′ reference point for each gene and define the distribution of the prior for the loading location *μ*_*L*_ to be a normal distribution centered at *z*_5_ (recommended) with width hyper-parameter *a*. Importantly, we also use *z*_5_ for the *μ*_*L*_′ prior, consistent with the bidirectional nature of loading and initiation. On the other hand, we found the actual dissociation location (*μ*_*T*_) is more variable gene-to-gene. Despite this variability, we can expect that it must occur somewhere downstream from the end of the mature transcript—for example, downstream of the PAS, the 5′end of the 3′UTR, OR the 3′end of the gene’s last exon. Therefore we assume this location is provided as the 3′ reference point *z*_3_—the upstream bound on the search for *μ*_*T*_. In this work we used the the 3′ end of the last exon for *z*_3_, with the exception of SOCS5 where the 5′ end of the 3′ UTR was employed. We make the further assumption that the dissociation location becomes decreasingly likely the farther downstream RNAP2 gets. Thus, we set the distribution of the prior for the termination location parameter (*μ*_*T*_) to be an exponential distribution, originating at *z*_3_ (recommended), with hyper-parameter *b* (for a diagram of these prior distributions see Supp. Fig. 8A). These prior positions, (*z*_5_, *z*_3_) are set by the user.

For the shape parameters that control the breadth of *LI* and *T* (*σ*_*L*_, *τ*_*I*_, *σ*_*T*_), we chose exponential prior distributions (user-defined). The prior distribution for the weight parameter (**w**) is a Dirichlet, as is convention for mixture models. Our default assumption is that all model components are equally likely, so we set all the *α* hyper-parameters for the Dirichlet priors equal to 1 (user-defined). The choices for all the priors are summarized in Eq. 13 (see diagrammatic depiction in Supp. Fig. 8B,C).

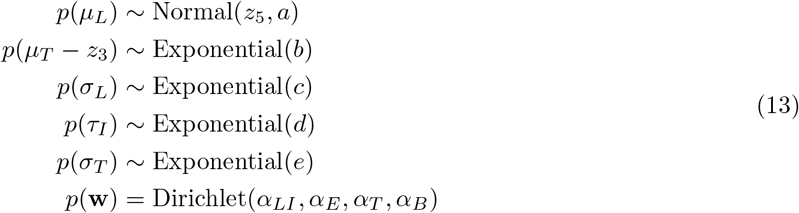

As discussed above, we provided a separate parameterization of the antisense-strand (Eq. 9). However, our software implementation provides flexibility in how the antisense-strand component of the model is handled, based on how the priors are specified in the input config file. There are three options: 1) tied parameters, where 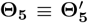 (similar to our previous model[23]); 2) independent parameters with equal priors, where 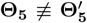 but 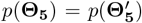; and 3) independent parameters with unequal priors, where 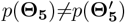. We recommend option 2, which can be interpreted as assuming the null hypothesis that the two strands have equivalent processes (asserted by the equivalent priors), but allowing the fit to independently adjust the parameters for the two strands, based on the data provided for fitting—i.e. priors for 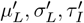 are equal to their sense-strand counterparts in Eq. 13. We demonstrate the impact of choosing option 1 or 2 by comparing their results in Fig. 3. Option 3 is provided for completeness and should only be used when there is *a priori* reason to believe that there is a systematic difference in *LI* between strands.

**Fig. 3:**
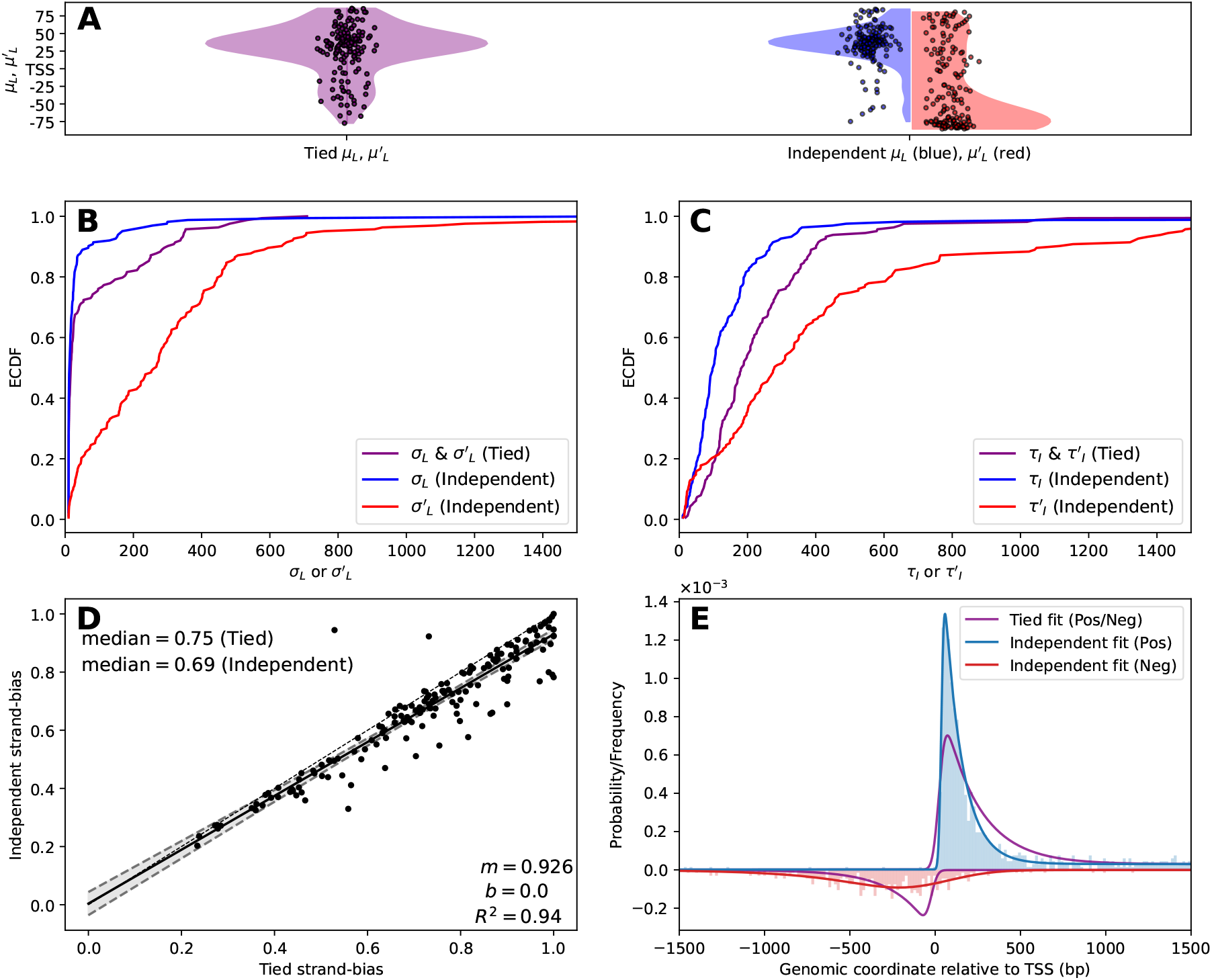
Modeling 5′ peaks independently uncovers distinct shapes between the strands. (A) Empirical cumulative distribution function (ECDF) of (B) loading position uncertainty parameters *σ*_*L*_, *σ*′_*L*_ and (C) characteristic initiation lengths *τ*_*L*_, *τ*′_*L*_ for the tied (purple) and independent (blue/red) scenario. (D) The calculated strand bias (Eq. 14) for the tied (x-axis) and independent (y-axis) scenarios. (E) Schematic of generated data from the median of 5′ parameters from the independent fitting scenario results, fit by the tied (purple) or independent (blue/red) scenario. Note that the independent scenario fits well to the data while the tied scenario cannot account for the differing strand profiles.

Note, our software allows the user to define the reference points (*z*_5_, *z*_3_) for each gene, the prior distributions for each (non-weight) parameter, and the values of the hyper-parameters for each prior distribution. The choice of distributions specified in Eq. 13 are the default recommendations and were used for all analysis herein. The following hyper-parameter values were used for all analysis: (*a, b, c, d, e*) = (1500, 500, 500, 10000, 500) (see Eq. 13). However, these values may not be optimal for all applications. As is typical with any Bayesian inference, tuning the hyper-parameter values may be necessary to optimize the fit results.

Our software implementation of the LIET model was written in Python 3 with the model-building and Bayesian inference being performed by the PyMC library (v5.6.1)([34]). The LIET model software implementation is available on GitHub at: github.com/Dowell-Lab/LIET.

### 2.3 Gene and sample selection

In order to showcase the capabilities of the LIET Model and draw conclusions from its fits, we needed a set of genes suitable for the model to which we can apply it and a range of high-quality data sets in which to fit those genes. The samples used in this paper were curated from a recently established nascent run-on sequencing database, DBNascent[17], which has been organized around sample metadata. We curated 152 high-quality PRO-seq datasets, generated from 24 different human cell lines, spanning 12 different tissue types (see Supp. Table 1 for samples). Most of these samples were in control conditions with the goal of analyzing how consistent basal termination was across cell-type (see Fig. 4). To evaluate the impact of a perturbation on the 3′ end dissociation position, we also included data from an Integrator knock-down experiment[25] and a heat shock experiment[35] (see Fig. 5).

**Fig. 4:**
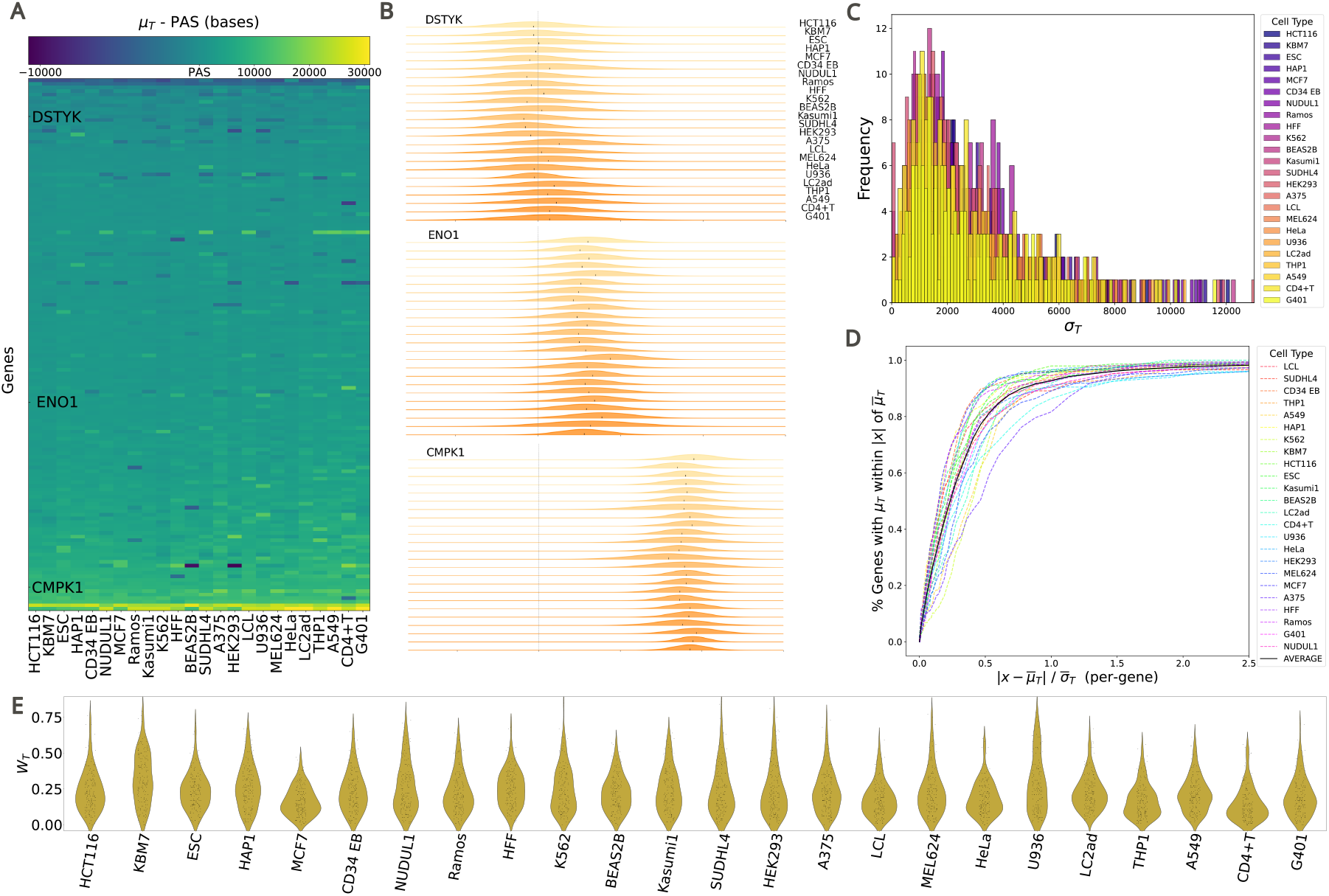
Termination parameters *μ*_*T*_ and *σ*_*T*_ are consistent across 24 cell types. (A) Heatmap of the distance between the PAS and *μ*_*T*_ per-gene, across 24 cell types. Genes (y-axis) are sorted on mean PAS - *μ*_*T*_ distance and cell types (x-axis) are sorted by the standard deviation of PAS - *μ*_*T*_ across genes. (B) Ridge plots of the dissociation distribution (*μ*_*T*_, *σ*_*T*_) across cell types for 3 genes: DSTYK (top), ENO1 (middle), and CMPK1 (bottom). Genes chosen to highlight the variation in *μ*_*T*_ (black ticks) positioning between genes. (C) Histogram of *σ*_*T*_ across cell types. (D) Cumulative distribution of distances of each inferred *μ*_*T*_ from the average dissociation distribution. Black line is the overall average across all cell types. (E). Violin plots of *w*_*T*_ across cell types.

As the model is designed to describe a single gene, we sought to identify a set of transcribed genes with no other transcription units within the fitting window, including no overlapping genes, enhancer RNAs, or lncRNAs. To arrive at our gene set, we first performed a number of computational pre-filters on the set of all known protein-coding genes in the human genome (from NCBI RefSeq transcript annotations, hg38, see Supplemental materials), eliminating those genes with other annotations within a set distance (10kb upstream and 30kb downstream) of the gene or transcription levels below a coverage threshold of 0.1x over the annotated gene body, averaged across all samples. These filters reduced the approximately 20,000 protein-coding genes down to approximately 1,400 candidate genes. These candidate genes were then manually inspected to identify cases of overlapping enhancer associated RNAs or other unannotated transcription units (see Supp. Sec. 4 for full details). We identified 163 genes spanning chromosomes 1–6 for subsequent testing. For a detailed description of the gene filtering and selection process see Supp. Sec. 4. The resulting set contained genes from approximately 1kb up to 200kb+ in length and of a range of different profile shapes and transcriptional levels. For the list of genes see the Supp Table 1.

Most published PRO-seq datasets are not sequenced deeply—typical gene coverage for a gene within these samples are well below 0.1x[17]—which adds to the difficulty of modeling transcription from these data with high precision. To ameliorate depth related issues arising during model validation, we created “meta-samples” by combining all technical and biological replicates for each cell-type. Some cell-types had many replicates (e.g. HCT116, HeLa, LCL) while others only had two (e.g. A549, NUDUL1, THP1). These meta-samples were then fit and analyzed for Fig. 3 and 4. For the cross-sample reproducibility analysis (Fig. 2) and perturbation analyses (Fig. 5), LIET was instead applied to the individual samples within that cell-type/condition. We found that the model maintains high accuracy across all parameters even when fitting to individual samples and low data coverage, but, as one would expect, the precision is impacted for some parameters (see Supp. Fig. 6 and 7).

**Fig. 5:**
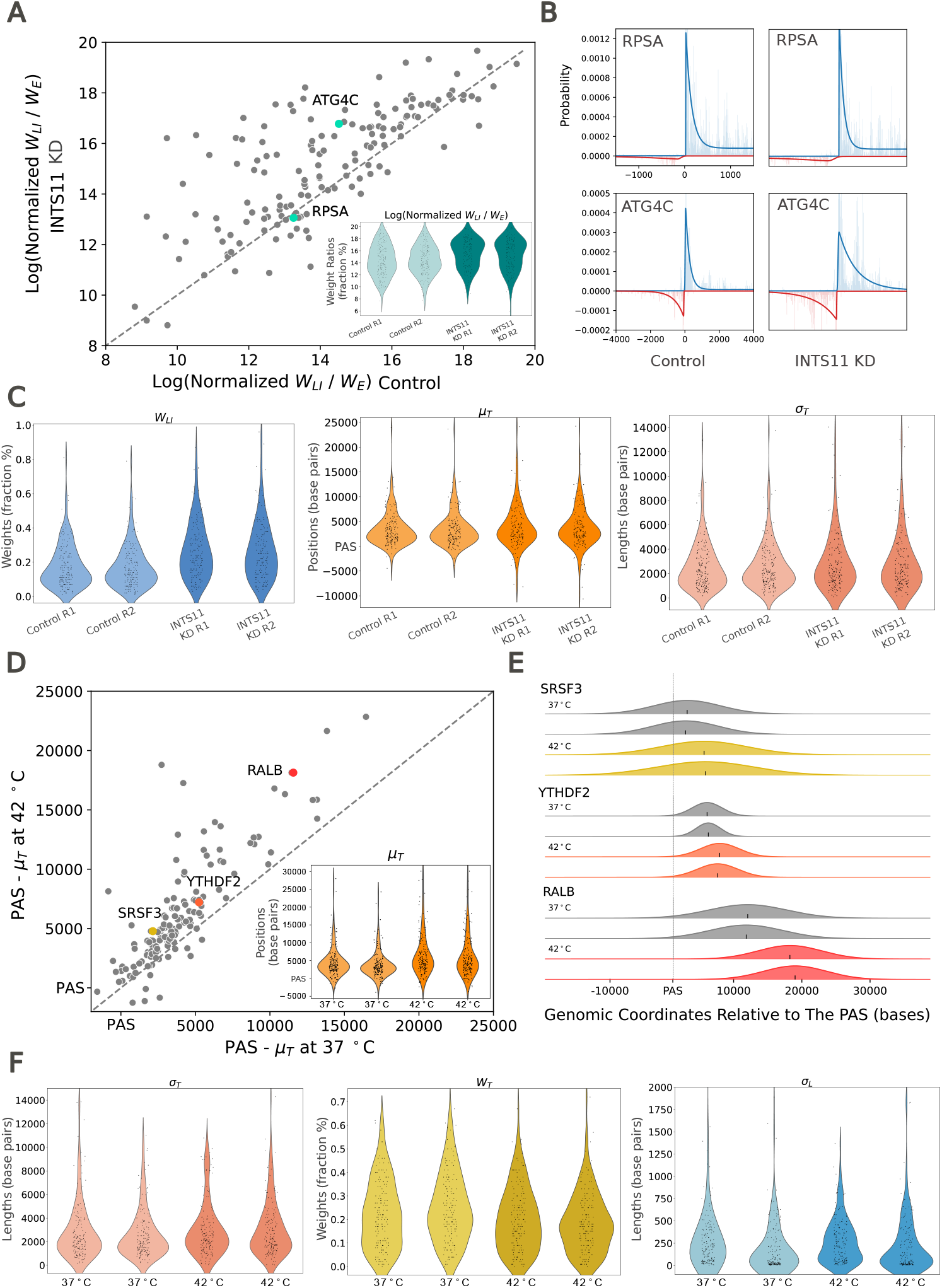
LIET model detects perturbation-induced changes in transcription. (A) Scatter and violin plot of the log of the averaged normalized pausing ratio (Eq. 15) across replicates for samples treated with and without an INTS11 knock-down (KD). Normalized pausing ratio increases at a subset of genes. (B) Bidirectional transcription at the 5’ end of RPSA and ATG4C with and without addition of the INTS11 KD. The pausing ratio for RPSA remains relatively unchanged upon treatment with the INTS11 KD, while ATG4C shows an increase in pausing ratio under the same conditions. (C) Violin plots showing control and INTS11 treated replicates across *w*_*LI*_, *μ*_*T*_, and *w*_*T*_. *w*_*LI*_ increases under exposure to the INTS11 KD, while termination-associated factors *μ*_*T*_, and *w*_*T*_ do not significantly change. (D) Scatter and violin plot of the average PAS - *μ*_*T*_ values across replicates for control and heat shocked samples. Heat-shock globally increases the distance between PAS and *μ*_*T*_ on a per-gene basis. (E) Modeled termination peak in SRSF3 (top), YTHDF2 (middle), and RALB (bottom). The position of *μ*_*T*_ is indicated by a black tick for each replicate, and the width of each peak is reflective of *s*_*T*_. *μ*_*T*_ at RALB shifts farther down stream when exposed to heat shock compared to SRSF3 and YTHDF2. (F) Violin plots of *s*_*T*_, *w*_*T*_, and *w*_*LI*_ in control and heat shocked conditions. *s*_*T*_ and *s*_*L*_ are not impacted by heat-shock whereas *w*_*T*_ decreases when exposed to heat-shock.

### 2.4 Data processing and representation

The LIET model describes the expected location of active RNAP2 instances at a snapshot in time. However, the details of the nascent run-on sequencing protocol influences the extent to which a read’s mapping location differs from the true location of a corresponding RNAP2 instance[7]. Details such as protocol selection (GRO-seq or PRO-seq), library preparation strategy, and read-length all complicate the biological interpretation of where RNAP2 was located. This raises the question of how to represent individual reads in the input to LIET. For example, a single read could be represented by its 5′end, 3′end, midpoint, or all positions along its length (full read). Using the full read effectively smooths the data but also artificially inflates the amount of data by the length of the read (multiple counting). Therefore, LIET assumes each read is represented by a single position. In general, it can be argued that selecting the 5′end of all reads provides the greatest fidelity on inferring the position of RNAP2 loading while the 3′end position of the read provides the greatest fidelity on the RNAP2 termination process.

For this work, we chose to represent the reads by the genomic coordinate of their 3′ end, as the *Termination* component of the model is of particular interest to us (given this is the first instance of modeling this process). Importantly, our decision influences the interpretation of the 5′ location parameters 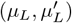 but not the shape or weight parameters. Specifically, location parameters at the 5′ end will be shifted downstream from the biological position of loading and initiation by a distance influenced by the expected length of RNAP2 run-on, the read length and other protocol details. In this work, we analyze a collection of data from numerous papers (see Supp. Table 1), each with distinct protocol details. Therefore we refrain from interpreting the 5′ positional parameters as direct readouts on the position of RNAP2 loading but rather focus on changes in this position between choices of priors (Fig. 3) or between samples (Fig. 5). Ultimately, future users of LIET must make their own decision on how to represent the data that best suits their application.

## 3 RESULTS

### 3.1 Design of a complete RNAP2 model

The activity of RNAP2, when transcribing a gene, is broken into four distinct stages: *Loading, Initiation, Elongation*, and *Termination*[1] (see Fig. 1A-C). The transcription process begins with the pre-initiation complex, which contains RNAP2, assembling at set locations in the genome (*Loading*). At genes, these assembly locations (parameter *μ*_*L*_) are near transcription start sites (TSS). Extensive CAGE data shows inherent variability in transcription start sites[36, 37], hence we model uncertainty in the loading position (parameter *σ*_*L*_). RNAP2 engages with one of the strands of DNA and subsequently transcribes a short distance in the 5′ 3′ direction and pauses (*Initiation*). Initiation is therefore the first step of transcription that produces RNA. At most genes, there is a second, upstream transcript (parameter *μ*′_*L*_ produced in the opposite direction[6, 38, 39]. These transcripts have been referred to as upstream antisense RNAs (uaRNA) or promoter upstream transcripts (PROMPTs)[38, 40, 41]. Thus we explicitly model two TSS in close proximity at every gene. We also assume there is an inherent uncertainty in the *Initiation* distance— that distance being more likely short than long (parameter *τ*_*I*_). After being released from its paused state, RNAP2 begins transcribing at an approximately constant rate in the 5′ → 3′ direction through the body of the gene (*Elongation*). The *Termination* stage of RNAP2 begins once RNAP2 transcribes the cleavage and polyadenlyation sequence (PAS)[42]. The PAS marks the end of the mature transcript, but RNAP2 proceeds well beyond the PAS, often for several kilobases or more[9, 42, 43]. RNAP2 appears to slow after the PAS before finally dissociating from the DNA[44, 45]. Therefore, we assume the position of dissociation (parameter *μ*_*T*_) is an unknown distance downstream of the PAS that is likely to vary between genes. We further assume variability (parameter *σ*_*T*_) in the duration of time between RNAP2’s slow-down and dissociation from DNA. Important for the LIET model, we assume a single instance of RNAP2 proceeds through these stages *sequentially* in time. Furthermore, because transcription proceeds only in one direction, we assume these stages must also be *spatially* sequential, for a single instance of RNAP2. Thus, if an instance of RNAP2 is undergoing *Elongation* it must be downstream of where it underwent *Loading/Initiation* and upstream of where it will undergo *Termination*). Given that nascent run-on techniques specifically target the RNA produced by RNAP at all steps of the transcription process, we describe each step mathematically based on the expected distribution of reads RNAP2 induces within nascent run-on data (Fig. 1; see Section 2.1 for full details). We implement our probabilistic mixture model and refer to the software as LIET.

The main objective of the LIET model is to accurately capture these processes and quantify their variation. A prototypical gene profile can be seen in Fig. 1D along with its LIET model fit (blue/red lines). This gene demonstrates the narrow, prominent sense-strand peak associated with *Loading* +*Initiation* near the TSS. It also has a much smaller antisense peak, indicating a significant sense-strand bias. There is a low and relatively uniform elongation region through the body of the gene. Lastly, a broader, less prominent pile up of reads is present downstream of the genes’ annotated end, which we associate with the dissociation process. The positive/negative strand residuals for the gene are plotted in Fig. 1E,F, and are representative of residuals observed across all genes analyzed. Across the entire profile, there is no systematic bias in the residuals and the average absolute residual is more than an order of magnitude smaller than the data signal, indicating the model is capturing all features of the data well.

### 3.2 Capturing the spectrum of transcription profiles

A cursory visualization of any deeply-sequenced nascent run-on sequencing dataset will highlight the extent of variation in the shapes of transcriptional profiles at protein-coding genes. The breadth and prominence of the 5′end bidirectional peaks, its sense/antisense strand-bias, the extent and depth of the elongation region, as well as the breadth and prominence of the dissociation peak all demonstrate significant variability across the gene landscape. Thus, we next sought to determine whether LIET is capable of accurately capturing this diversity. We show three example genes in Fig. 2A that differ in key elements of the gene profile. Gene CLIC4 is an example of a long gene (∼ 100kb) with an extreme sense-strand bias and large dissociation peak prominence. Gene UTP25 is an example of a medium length gene (∼ 30kb) with a significant antisense-strand bias at the 5′end and a dissociation peak of intermediate size located slightly upstream of the gene’s PAS. Gene MED18 is an example of a short gene (*<* 10kb) with a 5′ bidirectional region with coverage roughly balanced between strands. However, the two strands have fundamentally different shapes, with the antisense-strand *Loading* +*Initiation* distribution significantly more elongated than that on the sense-strand. Also, MED18 exhibits no clear dissociation peak. In place of a peak is a slowly decaying signal downstream of the PAS, which the model manages to fit by setting the termination peak weight (*w*_*T*_) close to zero. In short, these three genes demonstrate dramatic variation in their transcription profiles and the LIET model is sufficiently flexible to accurately capture them all.

We next sought to determine the extent to which LIET fits are influenced by experimental variability, as opposed to biological variation. To address this question, we independently fit the LIET model to each of the ten individual HCT116 samples (in control condition[28, 29, 30]—see Supp. Table 1 for SRRs) that comprise our HCT116 meta-sample (fitting the genes in our gene list) and then evaluated how well correlated all model parameters were for every pairwise sample comparison. Importantly, fitting individual samples lowers the statistical power of each fit, since the model is evaluating less data, but allows us to assess the sensitivity of LIET to the technical and biological variation that exists between samples. In Fig. 2B, we show the pairwise sample correlations for four model parameters across our gene list—two from the 5′end of the model (*w*_*LI*_ and *σ*_*L*_) and the two analogous parameters from the 3′end (*w*_*T*_ and *σ*_*T*_). All four model parameters show very high average correlation (*>* 0.74) within their respective publication (blocks along the diagonals). All correlations between publications are still moderately high (*>* 0.45), but the weaker correlations reflect differences in library preparation and sequencing protocol. In fact, the parameter correlation between experiments/publications is lower for the 5′end parameters than for those at the 3′end (e.g. A:C correlation for *σ*_*L*_, *σ*_*T*_ is 0.45, 0.8, in Fig. 2B), a result consistent with previous work showing the 5′end signal is particularly sensitive to protocol/library preparation[46]. A similar pattern is observed for all other model parameters—the average cross-sample correlation (averaged over all three experiments) for each parameter can be seen on the diagonal in Fig. 2C.

Another important aspect of model validity and interpretation is whether or not there exists spurious *cross-parameter* correlations (correlations between parameters that should be independent). To assess this, we calculated the correlation between every pair of parameters (averaged over all sample-pairs from the 10 individual HCT116 samples; Fig. 2C and Supp Fig. 9). Importantly, we expect some cross-parameter correlation in model weights *w*, due to their Dirichlet constraint (weights must sum to 1). Specifically, since the *LI* component overlaps with the *E* component in genomic coordinate space at the 5′end of the profile, the Dirichlet constraint indicates *w*_*LI*_ and *w*_*E*_ must be anti-correlated (a read in this overlap location would be assigned either to *LI* or *E*) and indeed that is the case (correlation *−*0.49). A similar argument holds for *w*_*E*_ and *w*_*T*_ at the 3′ end (*−*0.45). Also expected is the correlation between *μ*_*T*_ and *w*_*E*_—in general, the longer the *Elongation* region is, the more data it will contain and therefore the larger its weight (*w*_*E*_) will be. Likewise, the location parameters *μ*_*L*_ and *μ*_*T*_ define the approximate bounds of the *Elongation* region, giving an effective length (*μ*_*T*_ *− μ*_*L*_). Therefore *w*_*E*_ should be positively correlated with *μ*_*T*_ and negatively correlated with *μ*_*L*_. However, *μ*_*T*_ varies far more significantly gene-to-gene than *μ*_*L*_. Therefore the positive correlation with *μ*_*T*_ is more apparent than the negative correlation with *μ*_*L*_. Furthermore, due to the negative correlation of *w*_*E*_ with the other weights (*w*_*LI*_ and *w*_*T*_), *μ*_*T*_ would also be negatively correlated with them by the property of transitivity (coefficients −0.28 and −0.21). Importantly, all cross-parameter correlations (off diagonal values in Fig. 2C) are significantly lower than the parameters’ average auto-correlation (diagonal values), so we can reasonably assume the parameters are independent of one another during fitting.

### 3.3 Improved 5’end modeling captures strand differences

Transcription of the RNA upstream and antisense direction (the PROMPT) is generally concordant with the gene (i.e. both or neither are present[47]), but the rate at which RNAP2 engages with each strand (strand-bias) varies dramatically from gene to gene[23]. On the gene’s sense-strand, RNAP2 transitions from *Initiation* to *Elongation*, producing the pre-mRNA, whereas the PROMPT is rapidly degraded by the nuclear exosome complex. It is unclear to what extent the activity of RNAP2 at the PROMPT is distinct from that on the gene’s sense-strand and whether that difference could be regulatory. To address this question and better characterize RNAP2 activity between these two tightly spaced TSS’s, we used LIET’s robust framework to model the 5′ bidirectional region, allowing for independent parameterization of each strand. This is a distinct change from our earlier models of RNAP2[23] where the two strands’ 5′ peaks were assumed to arise from a single distribution (i.e. they were tied).

Therefore, we next sought to assess the impact of the two distinct 5′ end modeling options (tied or independent parameterization scenarios) on overall model fit. For this analysis we compared the two approaches on our HEK293 meta-sample—chosen to provide sample variety and examine a distinctly different cell-type. The “tied” parameterization 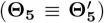 and the “independent” parameterization 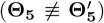 were run with the exact same priors—i.e. 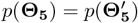. The results of this analysis are presented in Fig. 3, which shows how the 5^*1*^ parameter values differ under these two fitting scenarios.

First, we compare the mean loading position parameters *μ*_*L*_, *μ*′_*L*_ for the tied and independent fitting scenarios (Fig. 3A), measured relative to each gene’s annotated TSS. Unlike the tied approach, which results in a uni-modal distribution (purple in A), the independent fitting produces two distinct loading positions, one associated with each strand, and separated by a “footprint” of about 100bp. These distinct TSS’s have been observed in CAGE data[48] and the footprint has also been reported in previous work[18]. It is important to not be too critical of the absolute position of these two parameters relative to the annotated TSS’s, because we are utilizing a 3′ representation for input data which will shift the inferred position downstream a distance dependent on the read length and protocol (see Methods for detail).

We anecdotally observed differences in the shape of the distribution of *LI* between the sense and antisense-strands, so we wanted to determine if we could quantify this difference. To this end, we compared the empirical cumulative distribution function (ECDF) of loading position uncertainty parameters *σ*_*L*_, *σ*′_*L*_ for the two fitting scenarios (Fig. 3B). The tied scenario (purple) is intermediate to the other two (blue/red— from the independent scenario), due to the fact that it is a single parameter attempting to balance the data from both strands. It is more similar to the independent *σ*_*L*_ (blue), likely because there is typically more data on the sense-strand of a gene. In the independent scenario, the median values are 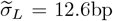 and 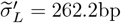 (tied median: 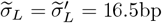). In other words, the typical loading uncertainty of RNAP2 is far lower on the mRNA-encoding strand. Next we compared the sense and antisense-strand characteristic initiation lengths *τ*_*I*_, *τ*′_*I*_ for the two fitting scenarios (Fig. 3C). Analogous to the distinction seen in the loading uncertainty, under the independent scenario, the median sense-strand initiation length 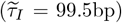 is significantly shorter than that for the antisense-strand 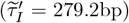. The dependent scenario result is again intermediate to the other two 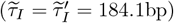.

How well the shape of *LI* (defined by *σ*_*L*_ and *τ*_*I*_) is inferred will impact how much of the data signal is allocated to the weights of these model components *w*_*L*_, *w*′_*L*_, from which the strand-bias is computed. In previous work, the strand-bias was treated as its own Bernoulli random variable[23]. In our case, we can compute strand-bias (denoted *π*) from the *LI* weight parameters and the total number of reads on each strand (*N*_+_, *N*_*−*_), in Eq. 14.

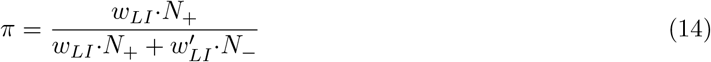

Strand-bias is defined on the interval [0, 1] and equals 1 for a given gene when all RNAP2 loads on the sense-strand. We compared the computed strand-bias under the two fitting scenarios using linear regression (Fig. 3D). The two scenarios are very well correlated (*R*^2^ = 0.94) but the independent scenario produces systematically lower strand-bias than the tied scenario, as indicated by the slope (*m* = 0.926) and a difference in medians of 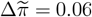.

In order to better interpret the differences in fitting the 5′ end peaks (both gene and PROMPT), we chose to run an illustrative simulation. Using the median parameter values from the independent scenario results in HEK293, we generated (simulated) data for a full gene profile to represent a prototypical gene. We then fit the simulated data under the tied and independent scenarios. The result of these fits (Fig. 3E) shows the difference in shape of the PROMPT (red data) from the gene sense-strand peak (blue data), consistent with real data (Figs. 1D and 2A). As expected, the independent fits (blue/red) are better at capturing the distinct shapes of each strand’s peak than the tied scenario (purple line) The tied scenario overestimates the breadth of the sense-strand peak while at the same time underestimating that of the antisense (PROMPT) peak, in an attempt to balance the shape of the two with a single set of shape parameters. Fitting the complimentary simulation—using median values from the tied scenario results—produces accurate results under both the tied and independent configurations (see Supp. Fig. 10). This highlights the flexibility in the independent modeling approach to fit a variety of shapes.

### 3.4 Modeling the 3’end identifies Termination consistency

Accurate quantification of the 3′end of gene transcription profiles has been a notoriously intransigent challenge[22] despite the fact that nascent run-on sequencing data has been available for over fifteen years[6]. The challenge arises in part because termination is the least well studied stage of RNAP2 activity[43]. From a modeling perspective, it is necessary to accurately identify the location of the end of the transcript profile as well as its shape—see the token examples in Fig. 1D and 2A. Unlike the 5′end where there are well annotated TSS, the 3′end of the transcription profile (i.e. the point of dissociation) is not annotated. Instead, the 3′ end annotation that most are familiar with refers to the end of the mature transcript, not the end of the region transcribed by RNAP2. Additionally, nascent run-on sequencing data sets appear to exhibit a gradual decay in signal at the 3′ end, making the identification of the last position of transcription highly sensitive to data quality and depth. To overcome these challenges, we include a dissociation-specific component in the LIET model (Fig. 1B) that captures the complete process of *Termination* (see Methods for full details).

Equipped with this new model, we sought to answer the questions: where does RNAP2 dissociate, and how consistent is it (in both location and shape) across cell-types? To this end, we calculate LIET fits on the meta-samples created from 24 different cell-types (Supp. Table 1) in basal/control conditions (see Section 2.3 for full details). We first examine the inferred position of dissociation, specifically asking whether the position of dissociation is reproducible across cell-types. With few exceptions, we see remarkable consistency across cell-types in the inferred position of dissociation relative to each gene’s PAS (Fig. 4A). Across all genes, the median distance from annotated PAS to *μ*_*T*_ location is 2478bp (upper/lower quartiles: *Q*_1_, *Q*_3_ = 1365bp, 4283bp). This is in contrast to the variation in *μ*_*T*_ between *genes* (examples in Fig. 4B). Interestingly, we find approximately 8% of gene fits where the inferred position of dissociation appears to be slightly upstream of the annotated PAS, but still downstream of the genes’ protein-coding sequences. We observe remarkable variation in the distance traveled by RNAP2 after the cleavage and polyadenlyation site. For example, RNAP2 travels roughly 10, 000bp farther downstream from the PAS at CMPK1 compared to DSTYK, the latter of which appears to terminate directly on top of the annotated PAS (Fig. 4B). Gene ENO1 demonstrates a termination position intermediate to the other two. Despite DSTYK terminating proximal to the PAS, it is more variable between cell-types and exhibits a greater spread (larger *σ*_*T*_) in the dissociation distribution, compared to the other two examples. Despite the variation some genes exhibit across cell-type, the variation observed in the *μ*_*T*_ position across genes is greater (e.g. comparing DSTYK and CMPK1). To further buttress the observed consistency, we ran a one-way, pairwise ANOVA on the distributions of PAS-to-*μ*_*T*_ distances (24 cell-types in Fig. 4A and Supp. Fig. 11) and determined that 99% of these pairwise comparisons (274/276) were not significantly different (p-adj *<* 0.05).

We then wondered if the width of the dissociation peak (*σ*_*T*_) differed at a given gene between celltypes. Generally, we observe that *σ*_*T*_ is also consistent (Fig. 4C) across cell-types. The median value for *σ*_*T*_ is 1926bp (upper/lower quartiles: *Q*_1_, *Q*_3_ = 1153bp, 3359bp). A one-way, pairwise ANOVA of the *σ*_*T*_ distributions identified no statistically significant differences between cell-types (p-adj *<* 0.05). Notably, an alternative way of considering the consistency of the dissociation position *μ*_*T*_ is to measure its variability relative to the width of the inferred dissociation distribution, quantified by *σ*_*T*_ (akin to a z-score). For example, consider an example gene from Fig. 4B from which we can compute its average *μ*_*T*_ and *σ*_*T*_ across cell-types. We can then assess for each cell-type if its individual *μ*_*T*_ is within *X* distance of the average *μ*_*T*_ (see x-axis in Fig. 4D). In this manner, we can quantify the consistency of *μ*_*T*_ across the cell-types for all genes in our list. We see that, on average, over 90% of a sample’s inferred *μ*_*T*_ values are within 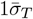 of each gene’s average *μ*_*T*_ (Fig. 4D)—further evidence for the consistency of the dissociation location. Ultimately, for those genes consistent across cell-type, the average *μ*_*T*_ values 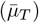 serve as a first annotation of the dissociation position, analogous to the TSS.

In contrast to the cell-type consistency of the dissociation location (*μ*_*T*_) and spread (*σ*_*T*_), the prominence of termination (*w*_*T*_) demonstrates significant cell-type specificity (Fig. 4E). Said another way, the fraction of reads present downstream of the cleavage and polyadenlyation site is, on average, low in some cell-types (examples: MCF7, LCL, HeLa) but quite large in others (examples: KBM7, K562, U936). The variability in the weight of the *Termination* component suggests this region may hold regulatory significance.

### 3.5 Detecting differential kinetics resulting from perturbation

One of the motivating goals of LIET was to leverage the model results towards a refined ability to identify gene specific patterns of differential RNAP2 activity, which manifest as changes to the shape of the nascent run-on sequencing data. Tools like DEseq[24] are designed to identify changes in gene expression levels on a per-gene basis. However, each gene is reduced to a single number—total reads over an interval—and therefore they lose details of how those reads are distributed. In contrast meta-gene analysis[27] calculates average profiles over a collection of genes, but struggle to identify which genes within the set have statistically different distributions of reads. The LIET model captures not only transcription-level information (**w, w**^′^) but also location 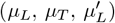 and shape information 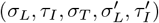, and therefore should be able to detect shape changes at the individual gene level. To test this aspect of the LIET model, we examine two previously published perturbation experiments.

Using a metagene approach, an Integrator knock-down (INTS11) was previously found to prevent RNAP2 from escaping from the 5^*′*^ pause state[25], increasing the magnitude of the 5^*′*^ peak of nascent run-on data. This is a compelling result, however, it cannot tell us whether the change is a global response or if it is instead dominated by a subset of genes (an inherent limitation to meta-gene analysis). Therefore, we first asked whether LIET can recover a similar result from the fit-inferred parameter values. For each gene, we plot the normalized pausing ratio (*ρ* ∝ *w*_*LI*_*/w*_*E*_; computed by Eq. 15—ratio of the *LI* and *E* weights, scaled by the widths of their respective distributions) in both control and Integrator knock-down conditions, averaging the ratio over the two replicates (Fig. 5A)—the replicates are reproducible, based on the distributions of the weight ratio seen in the inset and *w*_*LI*_ in Fig. 5C.

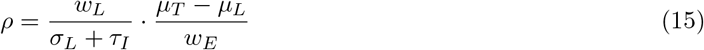

In Fig. 5A, we observe a clear increase in the pausing ratio at the majority of genes—given that most lay above the one-to-one line (dashed line)—indicating this knock-down is more or less a significant global perturbation (one-way pairwise ANOVA p-adj *<*0.05). However, the strength of this response varies dramatically gene-to-gene, with most genes showing a small response (ex. RPSA) while only a small number of genes demonstrate a large response (ex. ATG4C)—see Fig. 5B for the fits to these example genes. Given that Integrator is known to be critically important to the termination process at snRNAs and histone mRNAs[49], we next asked whether the 3^*′*^ end of our gene set (which contains no histone genes) were also altered in the knock-down. In this set of genes, we found no changes in the shape or location of the 3^*′*^ end (distributions for *μ*_*T*_ and *σ*_*T*_ in Fig. 5C—no significant differences in one-way pairwise ANOVA, p-adj = 0.66 and = 0.18, respectively).

It is also important to consider *how* the profile is actually changing: is the size of the 5^*′*^ peak simply increasing (larger *w*_*LI*_) or is there also a change to the *shape* of the peak? These two circumstances have different biological interpretations. If the only thing changing is the weight *w*_*LI*_ (see left plot in Fig. 5C), then one can say the nature of how RNAP2 undergoes loading and initiation is unchanged but RNAP2 fails to release from pausing in the knock-down. But if one sees an increase in the initiation length *τ*_*I*_ (ex. ATG4C in Fig. 5B), one could also infer that the fidelity of the initiation process has decreased. In summary, we see a decrease in *w*_*E*_ (complimentary to increase of *w*_*LI*_), an increase in *σ*_*L*_, and no changes in *μ*_*T*_ or *σ*_*T*_. Thus, LIET empowers us to tease apart the details of this perturbation. Plots of the distributions for all model parameter values (and their pairwise ANOVA results) for this experiment are in Supp. Fig. 12.

The second case study we employ focuses on heat shock, as this perturbation has been used to study run-through transcription[50]. We sought to determine whether LIET could identify changes in the position of RNAP2 dissociation in the run-through condition. For each gene, we plot the distance from the PAS to the location of *μ*_*T*_ obtained from the fits (the length PAS −*μ*_*T*_, averaged over replicates) for control (37°C) verses heat shock (42°C) conditions (Fig. 5D). Most genes lay above the one-to-one line, indicating most genes exhibit extended run-through (inset distributions indicates replicate reproducibility) in heat shock. The difference in distance traveled beyond the PAS (i.e. PAS −*μ*_*T*_) is significantly different between control and heat shock (one-way pairwise ANOVA p-adj*<* 0.05). Notably, the length of extended transcription varies gene-to-gene (distance of each point from diagonal line). For example, we highlight genes SRSF3 and YTHDF2 which show a minor shift in *μ*_*T*_ under heat shock compared to the much larger downstream shift of RALB (plots of their dissociation distributions, Fig. 5E). Remarkably, there was no significant impact on the shape of the dissociation peak (*σ*_*T*_ in Fig. 5F; non-significant in one-way pairwise ANOVA, p-adj= 0.43), suggesting that the dissociation process itself is only relocated and is not otherwise perturbed. Similarly, we see no significant differences in *μ*_*L*_, *σ*_*L*_, and *τ*_*I*_ under heat-shock (one-way pairwise ANOVA, p-adj= 0.69, 0.99, 0.79, respectively), indicating the 5^*′*^ processes are consistent, which refines a previous claim of no differences at the 5^*′*^ pausing region[51]. Interestingly, we do observe a minor but statistically significant redistribution of read signal—the weights *w*_*LI*_ and *w*_*B*_ increase while *w*_*E*_ and *w*_*T*_ decrease (p-adj*<* 0.05 for all). The weight changes may be explained by a minor global increase in RNAP2 recruitment and initiation under heat-shock, but more work is necessary to confirm this hypothesis. Plots of the distributions for all model parameter values (and their pairwise ANOVA results) for this experiment are in Supp. Fig. 13.

Ultimately, these two perturbations showcase the LIET model’s ability to detect changes in location, shape, and weight parameters on a gene-specific level. Thus, LIET proves itself to be a powerful tool in understanding the impact of perturbations on RNAP2 activity.

## 4 DISCUSSION

We present a new, probabilistic RNA polymerase II model that leverages the fact that each stage of RNAP2 activity induces unique distributions within nascent run-on data. The model is applied on a per-gene basis, providing parameter descriptions for every gene. Changes in parameters between conditions readily identify the impact of a perturbation not only on read levels (counts) but also in the shape and positioning of key RNAP2 activity. The model is also the first to capture the RNAP2 process of termination—specifically dissociation, which provides reproducible, data-driven annotations of the position of dissociation in both wild-type and stress conditions. The advancement in modeling capability provided by the LIET model, opens new avenues for exploring foundational regulatory mechanisms manifested in nascent run-on sequencing data.

Using the model, we find that the shape of the loading and initiation peak on the antisense-strand (the PROMPT) is inherently different than the peak on the sense-strand for the average gene. This can be seen in the dramatic differences observed in loading uncertainty *σ*_*L*_ vs. *σ*′_*L*_ and initiation length *τ*_*L*_ vs. *τ*′_*L*_ when the two strands are fit independent of one another. Our interpretation is that the pausing position of the PROMPT is less precise and further downstream of its loading location, relative to the gene-encoding strand. The PROMPT is unstable and terminates by a different mechanism than the gene[52], both of which may contribute to the observed difference in peak shape. Furthermore, we find that the typical strand-bias appears to be greater than 0.5 (more than 50% of RNAP2 are recruited to the sense-strand of a gene), with the median value being around 0.7. However, it is intriguingly variable gene-to-gene, with many genes possessing a strand-bias counter-intuitively below 0.5 (with some as low as 0.2). The extent to which strandbias is utilized by the cell as another regulatory mechanism is unclear. LIET provides a tool for subsequent refined studies on the transcription process at PROMPTs and how this process relates to gene regulation.

The inclusion of an explicit model of termination, focused on RNAP2 dissociation from DNA, is a major improvement from the LIET model over prior models. We capture the dissociation process and find that the position and shape is reproducible across replicates and cell-types for unperturbed cells. In fact, the shape of the dissociation peak (*σ*_*T*_) is largely unchanged even in stress conditions when the position of the dissociation peak (*μ*_*T*_) shifts dramatically downstream. What influences the relatively precise positioning in either scenario is unclear. While the positioning of the dissociation is reproducible at any given gene, the distance traveled after cleavage is highly variable between genes. Some genes exhibit dissociation positions (*μ*_*T*_) immediately proximal to the PAS point whereas at other genes RNAP2 travels more than 10kb downstream. Likewise the weight of the dissociation peak (*w*_*T*_) varies across cell-types, consistent with variable gene transcription levels in the region downstream of the gene. Since most nascent run-on sequencing analysis pipelines quantify genes based on the annotation, they fail to include the large fraction of reads associated with the termination process (quantified by *w*_*T*_). This can have implications for understanding patterns of differential transcription elsewhere, including in the body of the gene, as a failure to account for a large fraction of mapped reads can throw off some normalization strategies. It will be important in future work to more broadly characterize run-through transcription, as the LIET framework adds a degree of quantifiability and precision to the process that was previously lacking.

Our analysis here focused on a relatively small set (163) of isolated genes, as our focus was primarily on the accuracy and reproducibility of the model. The LIET model describes RNAP2 activity at a single gene. Consequently, the presence of overlapping transcripts, for example enhancers within introns[17], complicates application of the model more broadly. Consequently, our next goal for the model is to characterize its performance when the isolation assumption is violated. Three features of the model suggest it can be applied more broadly to non-isolated genes. First, the *Termination* component is Gaussian and occurs completely on the sense strand. This will help the model distinguish dissociation peaks from the characteristic bidirectional EMG shaped peaks present at RNAP2 loading and initiation—a shape also inherent to enhancer transcription—that may be proximal to a gene’s 3^*′*^end. Second, because LIET captures elongation as a mixture of the 5^*′*^ and 3^*′*^ components, enhancers—which are lowly transcribed—residing within introns may not strongly impact the model. Third, the model’s strand-specific background components help to account for data beyond the bounds of the transcription profile that cannot be attributed to the profile itself. Even in cases where the density of transcription is problematic, we may find ways of leveraging details of the underlying protocol differences or orthogonal data (for example H3K27ac ChIP) to dissect the contributions of individual transcripts.

We took a conservative approach to the conceptualization and derivation of the LIET model: we based the model components on the well-established stages of RNAP2 transcription. However, we have reason to believe that the model is biologically relevant to a broader range of types of transcribed loci. For example, though the details of transcription by RNA polymerase I (RNAP1) is known to be distinct from that of RNAP2, we believe the LIET model is sufficiently flexible to capture RNAP1 loci, the results of which could be used to quantify the contrast between the two. Furthermore, by systematically comparing LIET model fits of lncRNA (or any other class of RNAP2 transcripts) to that of protein-coding genes, we can quantify any distinction in RNAP2 activity. Thus, we intend to apply the LIET model to a greater variety of loci.

A few other improvements to the LIET software would extend its utility. First, other more common protocols, such as Pol II ChIP and metabolic labeling, provide similar information (but with unique data characteristics) to nascent run-on assays. Therefore adapting LIET to these datasets would broaden its use. Second, we will continue to find ways to make LIET more efficient, as currently it is accurate but not particularly fast. Here we side step the efficiency issue by simple parallelization, as each individual gene can be run without regard for the others. However, for whole genome analysis or application to hundreds of samples[17], we will likely need more direct software improvements such as rewriting time consuming portions into C or C++. Finally, a formal framework for assessing significance of differential parameters would both add statistical power and streamline the identification of changes in the face of perturbations.

## Supporting information

LIET_model_supplement

## Funding

This work was funded by the National Science Foundation under grants ABI1759949 and the National Institutes of Health grant GM125871 and HL156475. The NSF NRT Integrated Data Science Fellowship (award 2022138) from the Biofrontiers Institute (to GEFB and HAT) and the Curci Scholarship from the Shurl and Kay Curci Foundation (to HAT) also enabled some of this work.

## Competing interests

The authors declare that they have no competing interests.

## Authors’ contributions

JTS designed, derived and implemented the model/algorithm. RDD provided high-level design for the model and advised on all aspects of the project. GEFB and JTS curated the final gene list and performed the data analysis presented in the figures. GEFB, HAT, and JTS validated the algorithm on published data. HAT helped with software debugging and preliminary data analysis. RFS helped with sample curation and algorithm design. MAA advised on the biological interpretation of the model, experimental design, and performed the initial gene curation. JTS and RDD wrote the manuscript. All authors revised the manuscript.

## Acknowledgements

We thank Zach Maas and Samuel Hunter for contributions to discussion on the algorithm. We thank Jen Kugel for consultation on run-through transcription. We are also grateful to the BioFrontiers IT department for their support in building the database and maintaining the HPC system we relied on for all our analysis.

## Data Availability

Processed data and intermediate files can be found on Zenodo (10.5281/zenodo.10223322) and on the DBNascent website (nascent.colorado.edu).

## Code availability

All the code needed to run LIET is available at https://github.com/Dowell-Lab/LIET. The analysis to generate the figures is available within Jupyter Notebooks within the same repository. We also provide a guide for how to install and run LIET along with example inputs.

## Notes

https://github.com/Dowell-Lab/LIET

### Competing Interest Statement

The authors have declared no competing interest.

### Summary of Updates

Cleaned up language in the abstract and results section. Updated cross-referencing between the main text and the supplement. Included additional supplemental figures that support main figures 2--5. Cleaned up bibliography formatting.

