## Supplementary material for "LIET Model: Capturing the kinetics of RNA polymerase from loading to termination": LIET_model_supplement

### 1 Normalization schema for the Elongation distribution

The functional form chosen for the Elongation component of the LIET model (which we refer to as the "Elongation distribution" $E(z)$) is not an established probability distribution. It is the product of two expressions—the cumulative distribution function (CDF) of the exponentially modified Gaussian distribution (EMG, $5'$ end) and the survival function (SF) of the Gaussian distribution ($3'$ end)—whose individual integrals over $(-\infty, +\infty)$ are unbounded, but the integral of their product (Eq. S.1) can be shown to be bounded by a straightforward asymptotic analysis. Therefore, because this integral is bounded their product can serve as a valid probability distribution when scaled by the normalization factor. However, this function has no known anti-derivative and therefore a closed-form solution does not exist.

In this section, we present two normalization schema used for this function—one a straightforward but computationally inefficient numeric integration method and a second analytic integral solution that is highly accurate and computationally efficient. We universally recommend the use of the analytic solution, which we use for all analysis within the main paper. However, to validate the analytic solution and confirm its accuracy/precision, we compare it to the numeric integration method. We describe both in detail first.

**Disclaimer**: The notation used in this section is different than that used in the main text. We use different notation because 1) this section is a standalone mathematical exercise whose result has general utility beyond its role in the LIET model and 2) by using a simpler notation in this section, the math is easier to follow. In this section, we describe only the sense-strand Elongation component. Thus, the following is the correspondence between this section's notation and that in the main text.

$$\text{Parameter(supplement)} \qquad \text{LIET parameter(maintext)}$$

$$\mu \equiv \mu_L$$

$$\sigma \equiv \sigma_L$$

$$\tau \equiv \tau_I$$

$$\mu' \equiv \mu_T$$

$$\sigma' \equiv \sigma_T$$

First, the general problem of normalization consists of evaluating the integral given by Eq. S.1

$$A(\mathbf{\Theta}) = \int_{-\infty}^{+\infty} H(x \mid \mu, \sigma, \tau) \cdot G(x \mid \mu', \sigma') \, \mathrm{d}x \tag{S.1}$$

where $\mathbf{\Theta} = \{\mu, \sigma, \tau, \mu', \sigma'\}$ is the set of shape parameters for the (sense-strand) model and the functions $H(\cdot)$ and $G(\cdot)$ are the EMG ($5'$) CDF and Guassian ($3'$) SF respectively, defined as

$$H(x) = \Phi\left(\frac{x-\mu}{\sigma}\right) - \exp\left[-\frac{1}{\tau}\left(x - \mu - \frac{\sigma^2}{2\tau}\right)\right]\Phi\left(\frac{x-\mu-\sigma^2/\tau}{\sigma}\right) \tag{S.2}$$

$$G(x) = 1 - \Phi\left(\frac{x-\mu'}{\sigma'}\right) \tag{S.3}$$

where $\Phi(z)$ is the standard normal CDF (with $\phi(t)$ being the standard normal distribution) given by:

$$\Phi(z) = \int_{-\infty}^{z} \phi(t) \, \mathrm{d}t \tag{S.4}$$

$$\phi(t) = \frac{e^{-t^2/2}}{\sqrt{2\pi}} \tag{S.5}$$

The remainder of this section covers the two approaches used by the LIET software to evaluate Eq. S.1 during the process of model fitting.

##### 1.1 Numeric normalization method

Though the components of the LIET model are continuous functions—defined on $\mathbb{R}$ in the case of Loading, Elongation, and Termination and $\mathbb{R}^+$ in the case of Initiation—the data to which the model is applied is inherently discrete, defined on $\mathbb{N}^+$, since it represents genomic coordinates. Therefore, the maximum resolution needed for any numeric integration method is the integer scale. Thus, to compute $A$ numerically we need only choose a numeric integration range, evaluate the function at all integer points along this range, and calculate an approximate value for $A$ using a Reimann sum or trapezoidal rule.

Since the model is only valid at integer values, this method should be highly accurate and the degree of precision will be determined by the length of the integration range. We determine the integration range based on the variance of the distributions to which the functions $H(\cdot)$ and $G(\cdot)$ are related. The mean and variance of the EMG distribution are $\mu+\tau$ and $\sigma^2+\tau^2$ respectively.

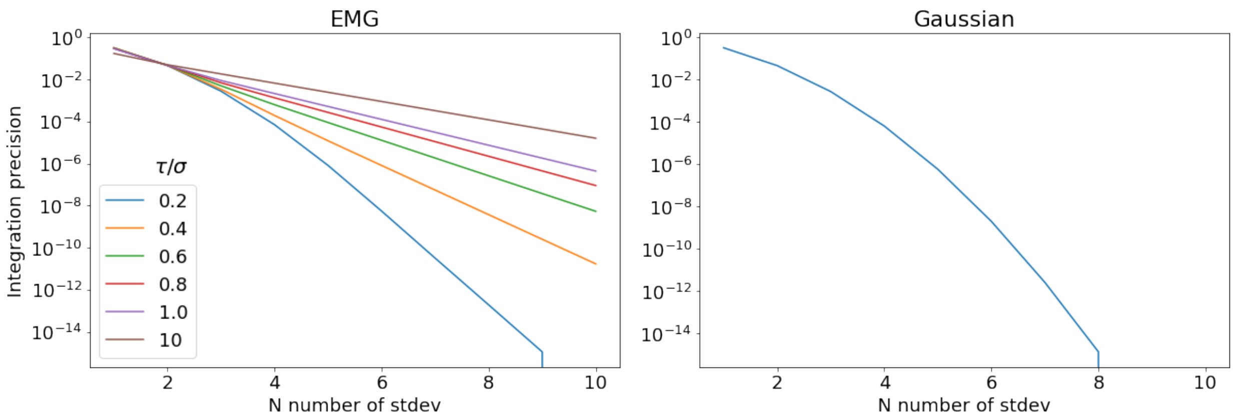


Supplementary Figure 1: The numeric integration precision for the EMG and Gaussian distributions as a function of the number (N) of standard deviations from the mean used to define the finite integral bounds. The integral bounds are defined as $mean \pm N \cdot stdev$. The precision for the two distributions are scale-invariant (i.e. the precision does not depend on the absolute value of the standard deviation), but for the EMG the relative proportion of $\tau$ to $\sigma$ has an impact on the precision of integration, especially for larger integration ranges (i.e. for larger N). However, for $N = 10$ the precision is $\leq 10^{-4}$ for all values of the shape parameters (given that $\tau/\sigma > 10$ is, effectively, not biologically feasible), which is sufficient precision for all practical model optimization instances.

For the Gaussian distribution, they are $\mu'$ and ${\sigma'}^2$. For the two individual distributions, Fig. **??** shows the precision of numeric integration using the number of standard deviations away from the mean as the integration bounds.

The precision of the numeric integration of the individual distributions is directly related to the precision of the product of their CDF and SF and determines our integration bounds for normalization of the Elongation distribution for any give set of values for $\mathbf{\Theta}$. Thus, we calculate the integration range $(z_{min}, z_{max})$ as follows:

$$z_{min} = \min\{\mu - n\sigma,\ \mu' - n\sigma'\} \tag{S.6}$$

$$z_{max} = \max\{\mu + n\sqrt{\sigma^2 + \tau^2},\ \mu' + n\sigma'\} \tag{S.7}$$

where $n$ is the integer number of standard deviations which determines the integration precision (as per Fig. **??**). In choosing the integration range in this manner, the precision is guaranteed to to have a fixed lower bound for the resulting Elongation distribution regardless of the parameter values. The normalization factor $A$ (Eq. S.1) is then computed as follows:

$$A(\mathbf{\Theta}) \sim \sum_{z_{min}}^{z_{max}} H(x \,|\, \mu, \sigma, \tau) \cdot G(x \,|\, \mu', \sigma') \tag{S.8}$$

One shortcoming to this approach is that for long genes (i.e. $\Delta = |\mu' - \mu|$ is large) with wide $5'$ or $3'$ transcriptional tails (i.e. $\sigma$ or $\sigma'$ are large) the integration range will be large, which is computationally prohibitive since this method requires the evaluation of the CDF and SF functions at every point along the range.

##### 1.2 Analytic solution normalization method

An alternative to the numeric method is to find an analytic solution to Eq. S.1 that is more easily computed. Here we demonstrate that this integral can be reduced to the evaluation of two standard normal CDF terms (i.e. there exists no closed-form solution for the $erf$ function), regardless of the parameter set. In other words computational evaluation of this solution is approximately constant in time, $O(1)$. What follows is a derivation of the the integral solution and demonstration of its limiting behavior.

The first step is to expand the CDF/SF product $H(x \mid \mu, \sigma, \tau) \cdot G(x \mid \mu', \sigma')$ into the sum of two terms. Each term can easily be shown to converge to zero as $x \to \pm\infty$ and therefore may be integrated separately. These two integrals are $I_1$ and $I_2$ below (Eq. S.9 and S.10).

$$A(\mathbf{\Theta}) = I_1 - I_2$$

$$I_1 = \int_{-\infty}^{+\infty} \Phi\left(\frac{x-\mu}{\sigma}\right)\left[1 - \Phi\left(\frac{x-\mu'}{\sigma'}\right)\right] \mathrm{d}x \tag{S.9}$$

$$I_2 = \int_{-\infty}^{+\infty} \exp\left[-\frac{1}{\tau}\left(x - \mu - \frac{\sigma^2}{2\tau}\right)\right] \Phi\left(\frac{x - \mu - \sigma^2/\tau}{\sigma}\right)\left[1 - \Phi\left(\frac{x-\mu'}{\sigma'}\right)\right] \mathrm{d}x \tag{S.10}$$

First we address integral $I_1$ (Eq. S.9). Apply a change of variables ($z = x - \mu$) and, using the Leibniz integral rule, we differentiate with respect to $\Delta$ where $\Delta = \mu' - \mu$.

$$\begin{aligned} \frac{\mathrm{d}I_1}{\mathrm{d}\Delta} &= \frac{\mathrm{d}}{\mathrm{d}\Delta} \int_{-\infty}^{+\infty} \Phi\left(\frac{z}{\sigma}\right)\left[1 - \Phi\left(\frac{z-\Delta}{\sigma'}\right)\right] \mathrm{d}z \\ &= \frac{1}{\sigma'} \int \Phi\left(\frac{z}{\sigma}\right) \phi\left(\frac{z-\Delta}{\sigma'}\right) \mathrm{d}z \end{aligned} \tag{S.11}$$

We can then reformulate the integral probabilistically: we define two independent standard normal random variables $Y$ and $Z$, and since $\Phi(\cdot)$ is the standard normal CDF we can re-express $\Phi\left(\frac{z}{\sigma}\right) \to \Pr\left(Z \leq \frac{z}{\sigma}\right)$. Applying a linear change of variables ($y = (z - \Delta)/\sigma'$), the integral becomes

$$= \int_{\mathbb{R}} \Pr\left(Z \leq \frac{\sigma' y + \Delta}{\sigma}\right) \phi(y)\, \mathrm{d}y$$

$$= \Pr\left(Z \leq \frac{\sigma' Y + \Delta}{\sigma}\right) = \Pr\left(Z - \frac{\sigma'}{\sigma} Y \leq \frac{\Delta}{\sigma}\right) \tag{S.12}$$

where we arrive at Eq. S.12 by the definition of the expectation $\mathbb{E}[f(Y)] = \int f(y)\phi(y)\mathrm{d}y$ and the use of the law of total expectation, such that $b \int \Pr(Z \leq a + by)\phi(y)\mathrm{d}y = \mathbb{E}[\Pr(Z \leq a + bY | Y)] = \Pr(Z \leq a + bY)$ Now we define the random variable $W = Z - \frac{\sigma'}{\sigma} Y$, and since $Y$ and $Z$ are both independent standard normal random variables, $W$ is also a normal random variable with a mean of zero and variance $\mathbb{V}\mathrm{ar}(W) = \mathbb{V}\mathrm{ar}(Z) + \frac{\sigma'^2}{\sigma^2} \mathbb{V}\mathrm{ar}(Y) = 1 + \frac{\sigma'^2}{\sigma^2}$. In other words, $W \sim \mathcal{N}\left(0, \sqrt{1 + (\sigma'/\sigma)^2}\right)$ which means $\frac{\mathrm{d}I_1}{\mathrm{d}\Delta}$ is equal to

$$= \Pr(W \leq \frac{\Delta}{\sigma}) = \Phi\left(\frac{\Delta/\sigma}{\sqrt{1 + (\sigma'/\sigma)^2}}\right) = \Phi\left(\frac{\Delta}{\sqrt{\sigma^2 + \sigma'^2}}\right) \tag{S.13}$$

In order to obtain the solution to the original integral $I_1$, this expression must be integrated with respect to $\Delta$. The function $\Phi(\cdot)$ has a well known antiderivative. Our final result is given by Eq. S.14.

$$I_1 = \int_{-\infty}^{\Delta} \Phi\left(\frac{\nu}{\sqrt{\sigma^2 + \sigma'^2}}\right) \mathrm{d}\nu$$

$$= \Delta \cdot \Phi\left(\frac{\Delta}{\sqrt{\sigma^2 + \sigma'^2}}\right) + \sqrt{\sigma^2 + \sigma'^2} \cdot \phi\left(\frac{\Delta}{\sqrt{\sigma^2 + \sigma'^2}}\right) \tag{S.14}$$

This solution cannot be further simplified since no elementary anti-derivative exists for the integrand of $\Phi(\cdot)$.

Next, we address integral $I_2$ (Eq. S.10). Here the added exponential term complicates the derivation, but we can again employ the Feynman integration method (differentiating with respect to $\Delta$ and after some simplifications integrating with respect to the same). We start with a change of variables ($x = \sigma z + \mu + \frac{\sigma^2}{\tau}$) prior to differentiating with respect to $\Delta$, to arrive at the following:

$$\frac{\mathrm{d}I_2}{\mathrm{d}\Delta} = \frac{\sigma}{\sigma'} \int_{-\infty}^{+\infty} \exp\left[-\frac{1}{\tau}\left(\sigma z + \frac{\sigma^2}{2\tau}\right)\right] \Phi(z)\, \phi\left(\frac{\sigma z - \Delta + \sigma^2/\tau}{\sigma'}\right) \mathrm{d}z \tag{S.15}$$

where, again, $\Delta = \mu' - \mu$. At this point, the integrand now contains two exponential terms (one linear in $z$ and the other, $\phi(\cdot)$, quadratic in $z$). We can then complete the square of their exponential terms to consolidate all dependence on $z$ into a single exponential term. In other words, we simplify the product of the two terms ($\exp[\cdot]\phi(\cdot)$) in the integrand of Eq. S.15 as follows (the details are left to the reader):

$$\begin{aligned} &\exp\left[-\frac{1}{\tau}\left(\sigma z + \frac{\sigma^2}{2\tau}\right)\right]\phi\left(\frac{\sigma z - \Delta + \sigma^2/\tau}{\sigma'}\right) \\ &= \phi\left[\frac{\sigma}{\sigma'}\left(z - \frac{\Delta}{\sigma} + \frac{\sigma^2 + \sigma'^2}{\sigma\tau}\right)\right]\exp\left[-\frac{\Delta}{\tau} + \frac{\sigma^2 + \sigma'^2}{2\tau^2}\right] \end{aligned} \tag{S.16}$$

Now we observe that by substituting Eq. S.16 back into Eq. S.15, the integral is of the same form as Eq. S.11, up to a scalar that is constant with respect to $z$, and therefore can be solved equivalently:

$$\begin{aligned} \frac{\mathrm{d}I_2}{\mathrm{d}\Delta} &= \exp\left(-\frac{\Delta}{\tau} + \frac{\sigma^2 + \sigma'^2}{2\tau^2}\right)\int_{-\infty}^{+\infty}\Phi(z)\,\phi\left[\frac{\left(z - \frac{\Delta}{\sigma} + \frac{\sigma^2+\sigma'^2}{\sigma\tau}\right)}{\sigma'/\sigma}\right]\frac{\mathrm{d}z}{\sigma'/\sigma} \\ &= e^{\Sigma/2\tau}\,e^{-\Delta/\tau}\cdot\Phi\left(\frac{\Delta - \Sigma}{\sqrt{\sigma^2 + \sigma'^2}}\right) \end{aligned} \tag{S.17}$$

where we have defined $\Sigma = \frac{\sigma^2+\sigma'^2}{\tau}$. Finally, in order to obtain our solution to $I_2$ we must now integrate Eq. S.17 with respect to $\Delta$

$$I_2 = e^{\Sigma/2\tau}\int_{-\infty}^{\Delta} e^{-\delta/\tau}\cdot\Phi\left(\frac{\delta - \Sigma}{\sqrt{\sigma^2 + \sigma'^2}}\right)\mathrm{d}\delta$$

to which we apply a change of variables $\nu = \delta - \Sigma$ in order to get the integral into the form of integral 101,000 in *A table of normal integrals*[1].

The resulting solution is given by:

$$
\begin{aligned}
I_2 =& e^{\Sigma/2\tau} e^{-\Sigma/\tau} \int_{-\infty}^{\Delta-\Sigma} e^{-\nu/\tau} \cdot \Phi\left(\frac{\nu}{\sqrt{\sigma^2+\sigma'^2}}\right) \mathrm{d}\nu \\
=& -\tau\, e^{-\Sigma/2\tau} \left[ e^{-\nu/\tau} \cdot \Phi\left(\frac{\nu}{\sqrt{\sigma^2+\sigma'^2}}\right) - e^{\Sigma/2\tau} \cdot \Phi\left(\frac{\nu+\Sigma}{\sqrt{\sigma^2+\sigma'^2}}\right)\right]_{-\infty}^{\Delta-\Sigma} \\
=& -\tau\, e^{-\Sigma/2\tau} \left( e^{-(\Delta-\Sigma)/\tau} \cdot \Phi\left(\frac{\Delta-\Sigma}{\sqrt{\sigma^2+\sigma'^2}}\right) - e^{\Sigma/2\tau} \cdot \Phi\left(\frac{\Delta}{\sqrt{\sigma^2+\sigma'^2}}\right)\right) \\
& + \tau\, e^{-\Sigma/2\tau} \left( \underbrace{e^{+\infty/\tau} \cdot \Phi(-\infty)}_{\to 0} - e^{\Sigma/2\tau} \cdot \underbrace{\Phi(-\infty)}_{\to 0} \right) \qquad \text{(S.18)} \\
=& \tau \left[ \Phi\left(\frac{\Delta}{\sqrt{\sigma^2+\sigma'^2}}\right) - e^{-\Delta/\tau} e^{\Sigma/2\tau} \cdot \Phi\left(\frac{\Delta-\Sigma}{\sqrt{\sigma^2+\sigma'^2}}\right)\right] \qquad \text{(S.19)}
\end{aligned}
$$

where the third term in Eq. S.18 is zero by L'Hôpital's rule and the fourth term is zero by $\Phi(-\infty) = 0$. As before, this solution cannot be further simplified due to the integral $\Phi(\cdot)$.

Lastly, by combining our result for $I_1$ and $I_2$, we arrive at our full solution for $A(\mathbf{\Theta})$—Eq. S.20,

$$
\begin{aligned}
A(\mathbf{\Theta}) =& \Delta \cdot \Phi\left(\frac{\Delta}{\sqrt{\sigma^2+\sigma'^2}}\right) + \sqrt{\sigma^2+\sigma'^2} \cdot \phi\left(\frac{\Delta}{\sqrt{\sigma^2+\sigma'^2}}\right) \\
& - \tau \left[ \Phi\left(\frac{\Delta}{\sqrt{\sigma^2+\sigma'^2}}\right) - e^{-\frac{1}{\tau}\left(\Delta-\frac{\Sigma}{2}\right)} \cdot \Phi\left(\frac{\Delta-\Sigma}{\sqrt{\sigma^2+\sigma'^2}}\right)\right] \qquad \text{(S.20)}
\end{aligned}
$$

where:

$$
\begin{cases}
\mathbf{\Theta} & = \{\mu, \sigma, \tau, \mu', \sigma'\} \\
\Delta & = \mu' - \mu > 0 \\
\Sigma & = \frac{\sigma^2+\sigma'^2}{\tau} \\
\sigma, \sigma', \tau & > 0
\end{cases}
$$

and therefore by combining Equations S.2, S.3, S.20 we arrive at the formal Elongation distribution:

$$
E(\boldsymbol{z} \mid \mathbf{\Theta}) = \frac{H(z \mid \mu, \sigma, \tau) \cdot G(z \mid \mu', \sigma')}{A(\mathbf{\Theta})} \qquad \text{(S.21)}
$$

###### 1.2.1 Limiting behavior of the analytic solution

In order to validate our analytic solution and confirm that it behaves as expected, we address some of the limiting behaviors of Eq. S.20.

First, it should be noted that the first two terms of Eq. S.20 (solution for $I_1$) would constitute the exact solution for the normalization factor if the Elongation distribution was instead a product of a *normal* CDF and normal SF. This is equivalent to $\tau = 0$ in our existing model of Elongation (Eq. S.21). Consider terms three and four in Eq. S.20 (the terms in the square bracket) when $\tau \neq 0$. If we factor out $\Phi(\Delta/\sqrt{\sigma^2 + \sigma'^2})$ (which we note is bounded on $(0.5, 1)$ given that $\Delta, \sigma, \sigma' > 0$), we can then focus on the expression

$$1 - e^{-\frac{1}{\tau}(\Delta - \frac{\Sigma}{2})} \cdot \frac{\Phi\left(\frac{\Delta - \Sigma}{\sqrt{\sigma^2+\sigma'^2}}\right)}{\Phi\left(\frac{\Delta}{\sqrt{\sigma^2+\sigma'^2}}\right)}$$

of which the second term can be shown to be bounded on $(0, 1)$ (asymptotically approaching 1 when $\tau \to \infty$ and 0 when $\sigma, \sigma' \to \infty$). Therefore we know integral $I_2$ is strictly positive. This implies $I_2$ acts as a "subtractive correction" to $I_1$ that accounts for the exponential initiation process, which comports with our intuition. In other words, the value of the normalization factor $A$ when $\tau = 0$ should always be less than that for $\tau > 0$.

Furthermore, we can consider the following special cases: $\sigma = \sigma' = \tau = 0$ and $\Delta = \tau = 0$. Intuitively, when $\sigma = \sigma' = \tau = 0$ the Elongation distribution should appear as uniform distribution of length $\Delta$ and therefore $A(\mathbf{\Theta}) = \Delta$. To show this we evaluate each of the terms in Eq. S.20 in this limit:

$$\lim_{\sigma,\sigma' \to 0} \Phi\left(\frac{\Delta}{\sqrt{\sigma^2 + \sigma'^2}}\right) = 1$$
$$\lim_{\sigma,\sigma' \to 0} \phi\left(\frac{\Delta}{\sqrt{\sigma^2 + \sigma'^2}}\right) = 0$$
$$\lim_{\sigma,\sigma',\tau \to 0} \tau\,[\;\cdots\;] = 0$$

So, Eq. S.20 under this limit is given by:

$$\lim_{\sigma,\sigma',\tau \to 0} A = \Delta \cdot (1) + (0 \cdot 0) - 0[(1) + e^{-\infty} \cdot (1)] = \Delta$$

Conversely, if we assume $\tau \neq 0$ and calculate the limit as $\sigma, \sigma' \to 0$, we would intuitively expect that the Elongation distribution would consist of a uniform distribution of length $\Delta$ with a "subtractive correction" (as discussed above) from an exponential distribution of characteristic length $\tau$. We see that our solution reduces to this form under said limit:

$$\lim_{\sigma,\sigma' \to 0} A(\mathbf{\Theta}) = \Delta \cdot (1) + (0 \cdot 0) - \tau[(1) + e^{-\Delta/\tau} \cdot (1)] = \Delta - \tau\left[1 - e^{-\Delta/\tau}\right]$$

This can be further simplified when the total transcription length is much longer than the initiation length (i.e. the case when $\Delta >> \tau$) to the expected result $A(\mathbf{\Theta}) = \Delta - \tau$.

Now to examine the other special case: when $\Delta = \tau = 0$. Although this is not a biologically relevant case for the purposes of fitting gene profiles since the approximate transcriptional length ($\Delta$) should never be zero in real world cases (i.e. it corresponds to a zero-length protein-coding gene), we should consider it for the purposes of confirming the limiting mathematical behavior of our solution for $A(\mathbf{\Theta})$.

In this case, the initial integral Eq. S.1 can be dramatically simplified to the following integral (which we denote as $A'$):

$$\begin{aligned} A' &= \int_{-\infty}^{+\infty} \Phi\left(\frac{x}{\sigma}\right) \cdot \left[1 - \Phi\left(\frac{x}{\sigma'}\right)\right] \mathrm{d}x \\ &= \int_{-\infty}^{+\infty} \Phi\left(\frac{x}{\sigma}\right) \cdot \Phi\left(-\frac{x}{\sigma'}\right) \mathrm{d}x \\ &= \int_{-\infty}^{+\infty} \Phi\left(y\right) \cdot \Phi\left(-\frac{y}{\sigma'/\sigma}\right) \sigma \mathrm{d}y \end{aligned}$$

which is in the form of integral 2,000.2 in *A table of normal integrals*[1] with $a = -\frac{\sigma}{\sigma'}$ ($< 0$) and thus the solution is as follows:

$$\begin{aligned} A' =& \sigma \left[x\Phi(x)\Phi(ax) + \phi(x)\Phi(ax) + \frac{1}{a}\Phi(x)\phi(ax) - \frac{\sqrt{a^2+1}}{a\sqrt{2\pi}}\Phi\left(x\sqrt{a^2+1}\right)\right]_{-\infty}^{+\infty} \\ =& \sigma \left[(+\infty)(1)(0) + (0)(0) + \frac{1}{a}(1)(0) - \frac{\sqrt{a^2+1}}{a\sqrt{2\pi}}(1)\right]_{x=+\infty} \\ &- \sigma \left[(-\infty)(0)(1) + (0)(1) + \frac{1}{a}(0)(0) - \frac{\sqrt{a^2+1}}{a\sqrt{2\pi}}(0)\right]_{x=-\infty} \end{aligned}$$

where the first term in each square bracket goes to zero by L'Hôpital's rule, so the result is

$$A' = \sigma \frac{\sqrt{(\sigma/\sigma')^2+1}}{(\sigma/\sigma')\sqrt{2\pi}} = \frac{\sqrt{\sigma^2+\sigma'^2}}{\sqrt{2\pi}}$$

This is equal to our general solution (Eq. S.20) for $A(\mathbf{\Theta})$ when $\Delta, \tau = 0$, where only the second term of the general solution is non-zero:

$$A(\mathbf{\Theta})|_{\Delta,\tau=0} = \sqrt{\sigma^2+\sigma'^2} \cdot \phi(0) = \sqrt{\sigma^2+\sigma'^2} \cdot \frac{1}{\sqrt{2\pi}}$$

##### 1.3 Precision of normalization schema

Though Eq. S.20 is indeed an exact solution and applicable in all parameter regimes, it is not necessarily machine-stable for all parameter values, therefore neither the numeric nor analytic normalization schema are of perfect precision. One may then ask how precise and equivalent to one another are the two methods? In order to explore this question, we first consider a representation of the analytic solution so that it is written in terms of proportionality between $\sigma, \sigma', \tau$ and $\Delta$. In other words, we define $\hat{\sigma} = \frac{\sigma}{\Delta}$, $\hat{\sigma}' = \frac{\sigma'}{\Delta}$, $\hat{\tau} = \frac{\tau}{\Delta}$. In this case, $\hat{\Sigma} = \frac{\Sigma}{\Delta}$. Therefore Eq. S.20 can be rewritten as follows:

$$\begin{aligned}\frac{A(\mathbf{\Theta})}{\Delta} =& \Phi\left(\frac{1}{\sqrt{\hat{\sigma}^2+\hat{\sigma}'^2}}\right) + \sqrt{\hat{\sigma}^2+\hat{\sigma}'^2}\cdot\phi\left(\frac{1}{\sqrt{\hat{\sigma}^2+\hat{\sigma}'^2}}\right) \\ &- \hat{\tau}\left[\Phi\left(\frac{1}{\sqrt{\hat{\sigma}^2+\hat{\sigma}'^2}}\right) - e^{-(1-\hat{\Sigma}/2)/\hat{\tau}}\cdot\Phi\left(\frac{1-\hat{\Sigma}}{\sqrt{\hat{\sigma}^2+\hat{\sigma}'^2}}\right)\right] \qquad \text{(S.22)}\end{aligned}$$

The fact that the entire solution can be written in terms of proportions means this normalization scheme is scale invariant—i.e. one can consider the behavior of the solution based only on the values of $\sigma, \sigma', \tau$ as a proportion of $\Delta$.

To test the precision and equivalence of our two schema across the parameter space, we evaluate the analytic solution (Eq. S.20) and the numeric solution (Eq. S.8) for a given set of values for the model parameters and calculate the percentage difference between the two values. Because the numeric method can be computed to a predetermined precision we use it as our reference value for the calculated percentage. We then did this over a logarithmic-scale grid of parameter values—$\hat{\sigma}, \hat{\sigma}', \hat{\tau} \in (0, 2]$.

Supp. Fig. S.2 shows the result of this comparison across the grid of values for $\sigma/\Delta$ and $\tau/\Delta$ (assuming $\sigma = \sigma'$). In general, we see extremely good agreement between the analytic normalization solution and the numeric normalization reference, with a maximum deviation of 0.0001%. The precision for the vast majority of the parameter landscape is limited by the data-resolution of $< 10^{-35}$. In Fig. S.2 there is one region of parameter space which has significantly lower precision than the rest—the region of large $\sigma/\Delta$ and low $\tau/\Delta$ (top left of figure). This arises due to the numeric instability of the last term in Eq. S.20 in which the exponential component $\exp(-(\Delta - \Sigma/2)/\tau)$ goes to infinity while the normal CDF component $\Phi(\cdots)$ approaches zero as

$\sigma, \sigma' \to \infty$, but their product is finite. Despite this instability, the precision of the analytic method in this parameter regime is more than sufficient for model fitting.

We can also assess the performance of the methods for asymmetric values of $\sigma, \sigma'$. In Supp. Fig. S.3 we show the percentage difference across a range of values for $\sigma$ and $\sigma'$ with $\tau = 0$. Effectively, this test isolates the precision of the first two terms in Eq. S.20. Here we see that precision is more or less uniform and high ($< 10^{-35}$) across the whole parameter space. This further supports the idea that the lower precision regions seen in Supp. Fig. S.2 are caused solely by the last term in Eq. S.20. Ultimately, both normalization schema show a high degree of precision and reproducibility across the full parameter space, making them suitable substitutes for the purposes of model fitting.

##### 1.4 Benchmarking normalization schema

To assess the compute time required for the two normalization methods, we timed repeated runs of the two methods on the parameter grid used for Fig. S.2 which contains a range of different parameter configurations. This was done on a single core with 4GB of memory. The numeric method took $481 \pm 3$s while the analytic method took $319 \pm 10$ms to compute the entire grid ($20 \times 20 = 400$ evaluations). Therefore, on average, the analytic method is approximately $10^3$ faster for realistic computing scenarios.

| |
|---|
| DISCLAIMER: As discussed at the start of Sec. 1, the notation used in this section is different than that for the LIET model in the main text. See the start of the section for parameter correspondence. |

#### 2 Uncertainty characterization and model validation

Here we demonstrate the model performance and parameter uncertainty of the LIET model by fitting to simulated data. These independent and identically distributed (*i.i.d.*) simulations demonstrate the relationship between the posterior widths, which provide estimates of uncertainty for parameter estimates, and the true variance of best fit parameter values (Supp. Fig. 4 and 5), thereby allowing us to contextualize the fit quality directly from the posterior widths. Furthermore, fits to these simulations serve as model validation

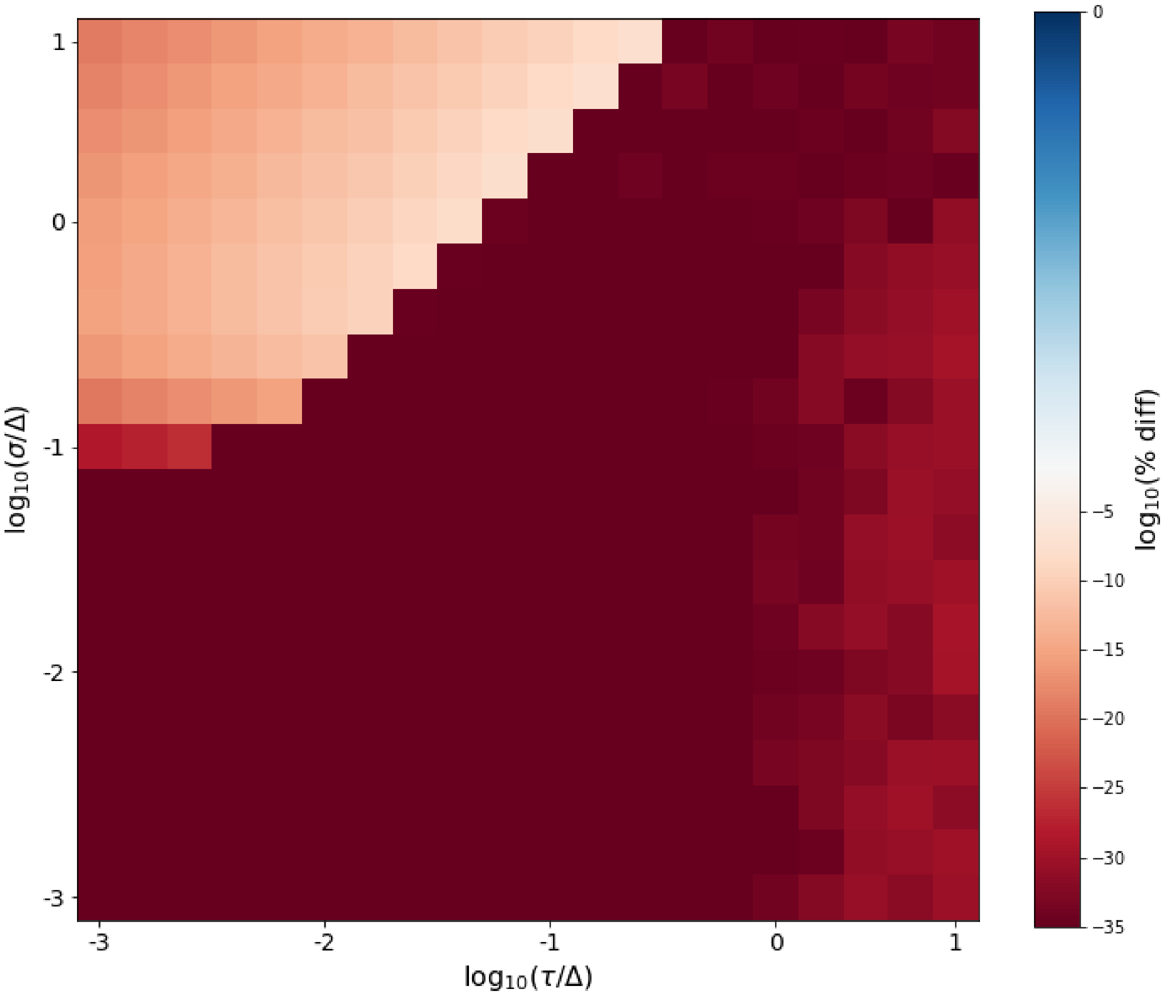


Supplementary Figure 2: Normalization precision for $\sigma, \sigma', \tau, > 0$: This heat map shows the fractional percentage difference between our two normalization schema across a logarithmic grid of parameter space, where we have set $\sigma = \sigma'$. In general, the two methods show extremely good agreement across all parameter values. The lowest precision regime is when $\sigma$ is relatively large and $\tau$ is small, however the worst precision is still 0.0001%, which is more than sufficient for the purposes of model fitting. Within the most biologically relevant region ($\sigma/\Delta, \tau/\Delta < 10^{-1}$), the two methods show effectively perfect agreement.

by demonstrating that model fitting recapitulates the correct ground-truth values for each model parameter (within the correctly quantified uncertainty) across a wide range of data qualities and levels of signal (Supp. Fig. 6 and 7). To perform the simulations, we set the gene size to reasonable parameter values representative of a prototypical PRO-seq gene profile. The parameter values used were as follows: gene length = 20kb, with the fitting window padded by 10kb upstream and 30kb downstream (i.e. size of the total fitting window = 60kb); shape parameters ($\mu_L = 0$, $\sigma_L = 1000$, $\tau_I = 500$, $\mu_T = 25000$,

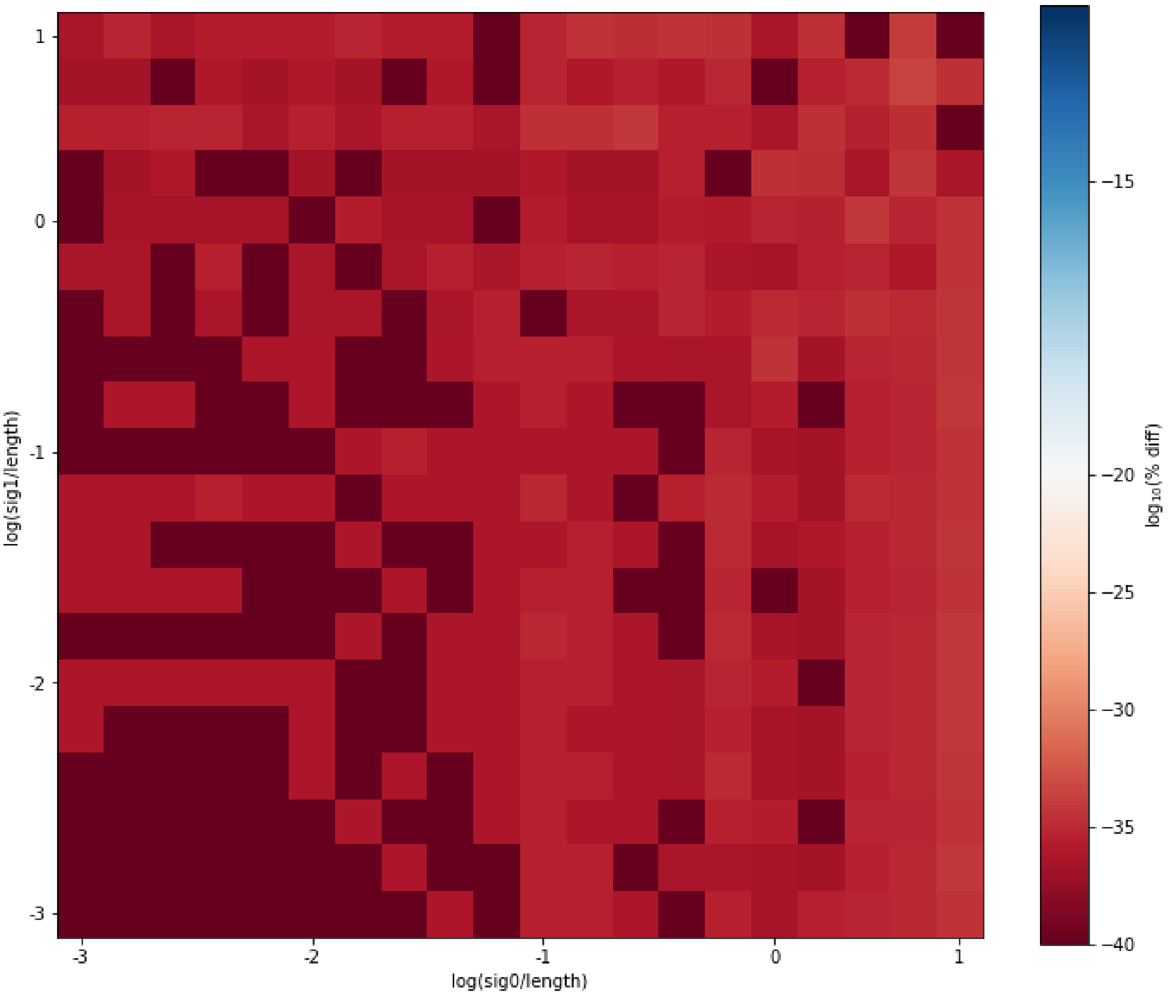


Supplementary Figure 3: Normalization precision for $\tau = 0$: This heat map shows the fractional percentage difference between our two normalization schema across a logarithmic grid of parameter space as a function of $\sigma$ and $\sigma'$ (we have set $\tau = 0$ but $\sigma$ and $\sigma'$ vary independently of one another). Given that $\tau = 0$, this test focuses specifically on only the first two terms in Eq. S.20. Furthermore, this tests the performance of the two normalization scheme for asymmetric configurations of the Elongation distribution. It is clear from the heat map that the two methods show effectively perfect agreement across the full range of parameter values, with the smallest precision being $\sim 10^{-33}$. Any variation seen in the heat map is likely due to the precision inherent to the float data-representation.

$\sigma_T = 2500$); and starting values for model component weights ($w_{LI} = 0.1$, $w_E = 0.7$, $w_T = 0.2$, $w_B = 0.0$). The anti-sense parameters were set equal to those of their sense strand counterparts. For the background/coverage assessment (Supp. Fig. 6 and 7), a range of total coverage and background weight values were used for the simulations, in order to demonstrate how the model fitting might perform over a range of data qualities. The total coverage ranged from 0.1x $\rightarrow$ 1.5x (a range comparable to or less than that of typical

PRO-seq data sets[2]). The background weights ranged from $w_B = 0.0 \rightarrow 0.5$. For $w_B > 0$, the remaining model component weights were uniformly scaled down from their starting values (see 'Reference' lines in Supp. Fig. 7).

Fitting to the simulated data was done using meanfield ADVI with an Adamax optimizer and 50,000 fitting iterations (implementation from `PyMC 5.6.1`). The simulation code can be found at the following GitHub repo https://github.com/Dowell-Lab/LIET.

##### 2.1 Calibrating the posterior uncertainty

One benefit to using Bayesian modeling methods is that the fitting process produces posterior distributions that serve both as parameter estimates and uncertainties associated with those estimates. However, depending on the fitting methods that are used, the uncertainties derived from the posterior distributions can be systematically biased. This is the case for most Variational Inference fitting methods, which systematically underestimate the uncertainties on parameter estimates[3]. For this reason, it is necessary to compare the uncertainty estimates obtained from the posterior to the true variation/uncertainty in the best estimates of the parameters if one wants to use the posterior to accurately estimate parameter uncertainty. To do this in a controlled fashion, it is helpful to use simulation to precisely define the model's ground truth and the signal-to-noise of the data (i.e. data depth relative to background).

We performed a number of *i.i.d.* simulations (as described in the previous section) but with varying levels of total coverage (range $10^4 \rightarrow 10^5$ reads). For each coverage value, we generated 10 simulations. Focusing on the termination component of the model (the portion of the profile that is the most difficult to model and that the LIET is the first to capture), we obtain the best fit parameter estimates for $\mu_T$ and $\sigma_T$ and uncertainties on those estimates from the means and standard deviations from the parameters' posteriors, respectively. These are plotted in the top two panels of Supp. Fig. 4 and 5 respectively. In the top panel of both figures, we see the average posterior width ($\bar{\sigma}_{post}$) decreases with increasing read coverage, suggesting an increasing confidence in our parameter estimates. However, looking at the middle panels of these two figures we see the true variation in the parameter best estimates (scatter of red points) are systematically greater across the full range of coverage values and do not decrease significantly as compared to the posterior widths (grey bands). This indicates the posterior widths are underestimates of the uncertainties on the parameter estimates. To quantify the disparity, we calculated the difference of each simulation's parameter estimate from the known ground truth value for the simulation (absolute value) and took the ratio of it to the simulation's posterior standard deviation. Effectively, this ratio measures how far off the posterior width is from the true deviation of the estimate. Looking at the distribution of this ratio for all simulations allows us to choose a statistically representative value to use for estimating true model uncertainty from our posterior distributions. Choosing a value between the upper quartile and the maximum ratio value excluding outliers would serve as a conservative yet reasonable estimate for this ratio. Based on Supp. Fig. 4 and 5, this would mean choosing a value for the ratio between 3 and 6 times the $\sigma_{post}$. As a conservative rule of thumb, we assume $|\widehat{\mu_T} - \mu_T|/\sigma_{post} = 3$. Therefore, the uncertainty for a given parameter estimate is assumed to be $3\sigma_{post}$.

##### 2.2 Model performance vs data quality

The *i.i.d.* simulations in Supp. Fig. 6 and 7 demonstrate the model performance for all shape parameters and model component weights as a function of data quality. In this case, "data quality" is quantified by total read coverage as well as the proportion of those reads assigned to background (as described at the start of this section). In general, the expectation for model performance is that as total read coverage goes down (i.e. there's less data) and/or the proportion of reads that are a part background goes up (i.e. signal-to-noise decreases) then precision on parameter estimates should get worse (i.e. parameter estimate uncertainty increases). This is what we observe in Supp. Fig. 6—the variability in the simulations increases for all parameters. Additionally, in these circumstances the accuracy of the parameter estimates should not be effected. In other words, there should be no systematic bias in the parameter estimates. In fact, this is what we observe in Supp. Fig. 6 and 7, where the average of the simulations are centered on the ground truth parameter values.

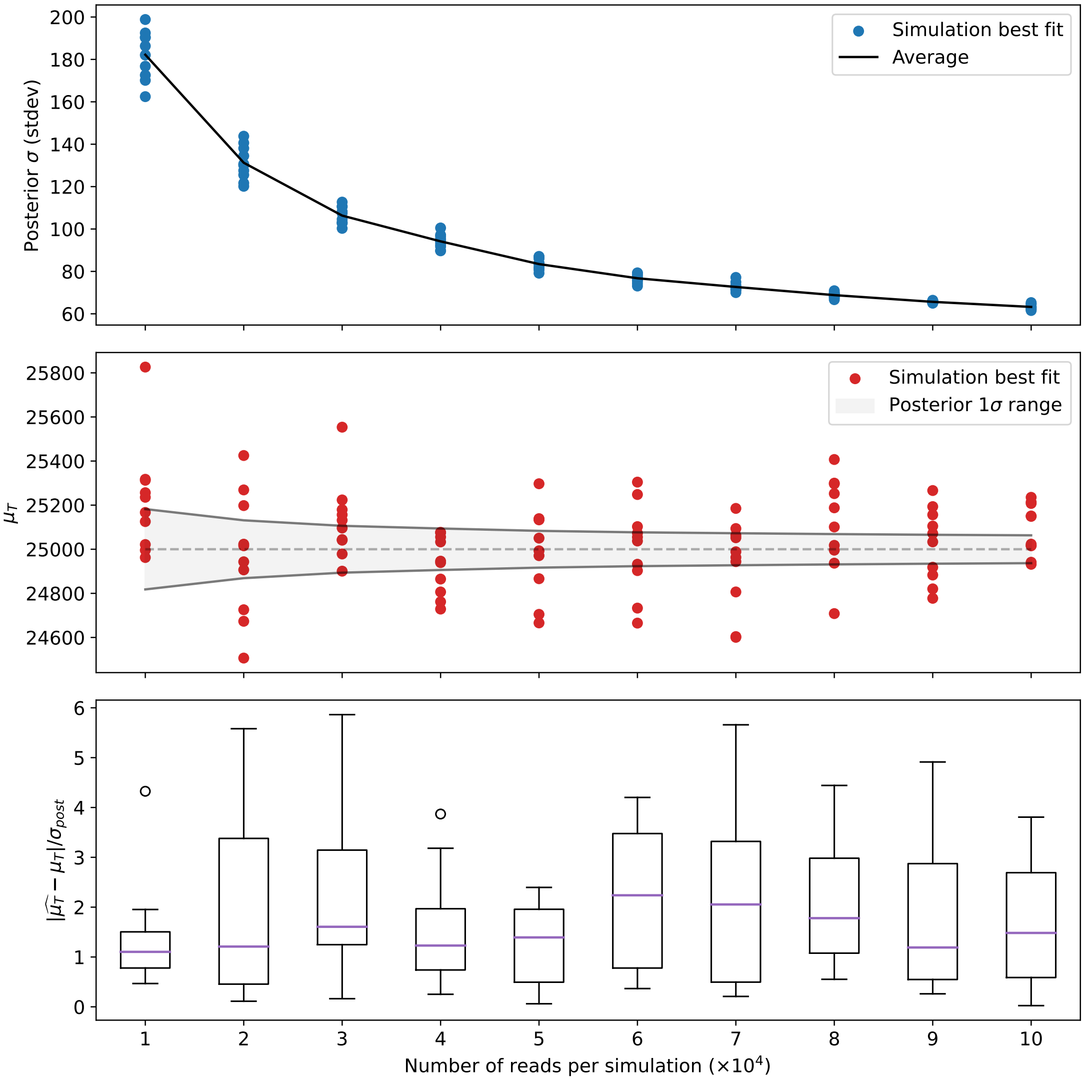


Supplementary Figure 4: Error characteristics for $\mu_T$. Using simulated data for a typical gene profile, we demonstrate the relationship between the posterior distribution widths and the error on the best fit parameter value—i.e. deviation of posterior means from ground truth values. This error arises from the stochasticity of finite data sampling (in simulated or real data). **Top**: Scatter plot of the standard deviation of $\mu_T$ posterior distributions (blue dots) from fits to multiple *i.i.d.* gene simulations, as a function of sampling depth (i.e. number of simulated reads). Black line indicates the average for the ten simulations at each sampling depth. Note, increasing sampling depth results in a narrower posteriors, indicating greater confidence in "best fit" value. **Middle**: Comparison of the best fit values for $\mu_T$ (red points—i.e. the means of the posterior distributions, $\widehat{\mu_T}$) to the $2 \cdot \bar{\sigma}_{post}$ width (black line, same as that in the Top plot) about the ground truth $\mu_T$ value (dashed gray line) for the same simulations. **Bottom**: The ratio of the deviation of best fit value (red points in Middle) from ground truth ($|\widehat{\mu_T} - \mu_T|$) to posterior distribution width ($\bar{\sigma}_{post}$—blue in Top) for each simulation, versus depth. Note: for all depths the median ratio is greater than 1.

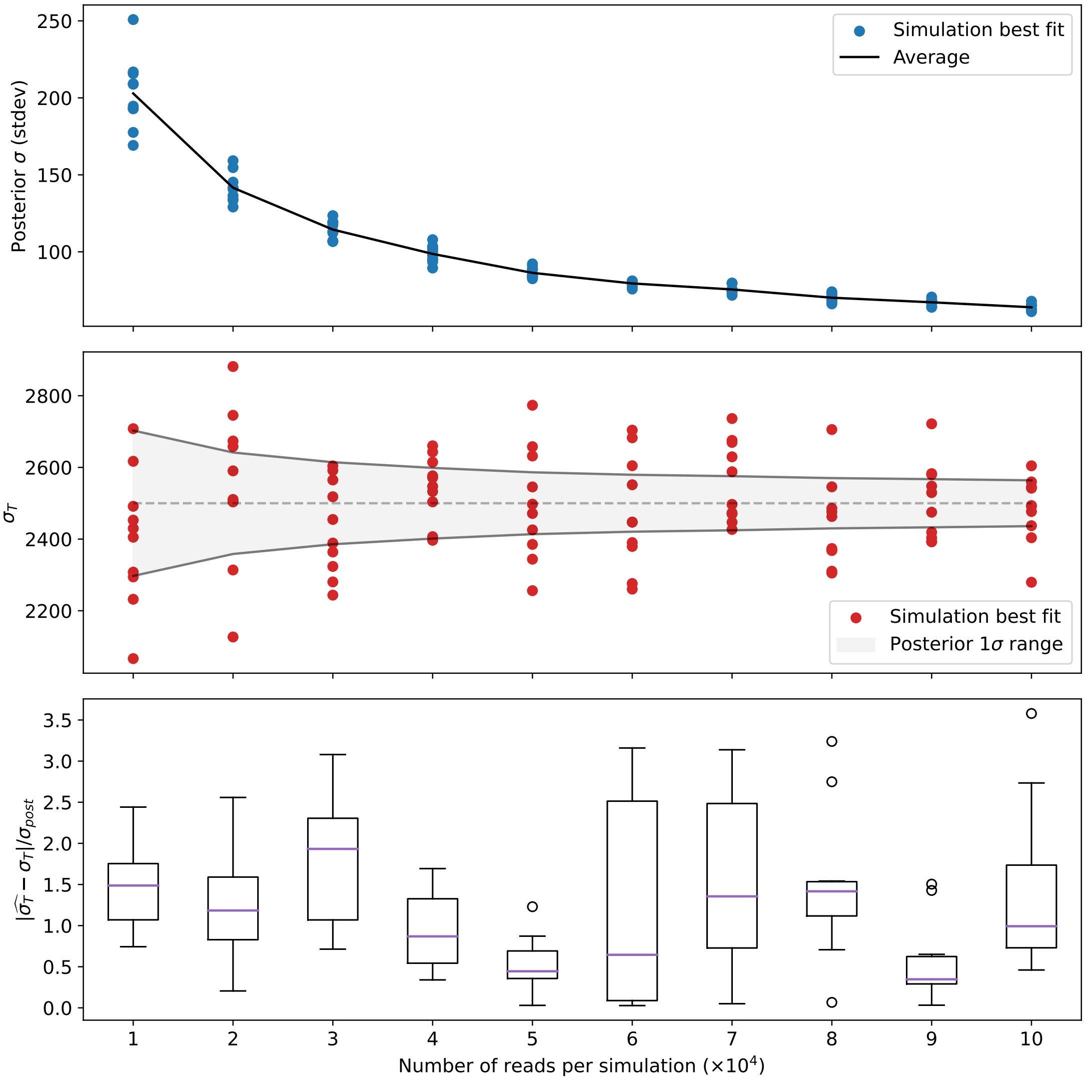


Supplementary Figure 5: Error characteristics for $\sigma_T$. Demonstration of the relationship between the posterior distribution widths and error on the best fit parameter value—i.e. deviation of posterior distribution means from ground truth values—due to the stochasticity in simulated data. Figure structured equivalent to Supp. Fig. 4 except results pertain to parameter $\sigma_T$—they were generated from the same simulations. When performing Bayesian inference, the breadth of the posterior distribution is the only measure of model parameter uncertainty and for variational inference methods this is a biased underestimate of the real parameter uncertainty. The error characteristics in this and Supp. Fig. 4 are useful in interpreting the realistic degree of uncertainty for a given parameter's best fit value. For example, the average posterior stdev $\bar{\sigma}_{post}$ could be scaled by a factor to bring it in line with the upper limits on the best fit value's deviation from ground truth (i.e. $3 - 6\bar{\sigma}_{post}$), and this value could serve as the parameter uncertainty. The simulations for this and Supp. Fig. 4 used the following ground truth parameter values: $\mu_L = 0$, $\sigma_L = 1000$, $\tau_I = 500$, $\mu_T = 25000$, $\sigma_T = 2500$.

### 3 Model parameter interpretations

This section elucidates some of the biological meanings/interpretations of the model's parameters.

#### 3.1 Background weights ($w_B$ and $w'_B$)

Because the model has differing numbers of mixture components between the two strands, this changes the interpretation of the component weighting on each of the two stands. Specifically, equal weighting to the background component of the mixture does not constitute equivalent absolute levels of background between the two strands. In other words, if $(w_B, w'_B)$ represent the sense and anti-sense strand background read proportions respectively and $(N_B, N'_B)$ represent the absolute number of reads within the fitting window assigned to background on the sense and anti-sense strands respectively, then $w_B {\equiv} w'_B \not\Rightarrow N_B {\equiv} N'_B$. However, the intuitive interpretation for "equivalent background" between the two strands is when the absolute number of background-assigned reads is equal between the two strands, i.e. $N_B = N'_B$. Here we show how $w_B$ and $w'_B$ are mathematically related under this circumstance. Furthermore, we can also account for the impact of strand-bias on this relationship.

Strand bias (which we label $\pi$—Eq. 14 in the main text) is the proportion of polymerase implicated in loading on the sense strand (thus, the proportion loading on the anti-sense strand is given by $1-\pi$). The total number of reads on the sense strand are given by $N = N_{LI} + N_E + N_T + N_B$. The total reads on the anti-sense strand are given by $N' = N'_{LI} + N'_B$. The strand bias relates $N_{LI}$ and $N'_{LI}$ as follows: $N_{LI}/N'_{LI} = \pi/(1-\pi)$. Therefore, if we consider the condition of "equal background" as defined in the previous paragraph, we find that the relationship between $w_B$ and $w'_B$ also depend on the strand bias and the proportion of reads assigned to Loading/Initiation on the sense strand, i.e. $w_{LI}$. This can be shown as follows:

$$
\begin{aligned}
N' &= \frac{1-\pi}{\pi} w_{LI} N + w_B N \\
\Rightarrow N \frac{w_B}{w'_B} &= N \left( \frac{1-\pi}{\pi} w_{LI} + w_B \right) \\
\Rightarrow w'_B &= \frac{\pi w_B}{(1-\pi) w_{LI} + \pi w_B}
\end{aligned}
\tag{S.23}
$$

Here, in the first equation we have used the strand bias relationship for the first term and the definition of "equivalent background" for the second term. It is clear from this relationship that direct comparison of model background weights for each of the two strands is not possible. In practice, it is more useful to consider the background in terms of the total number of reads present on each of the two strands and the background weight: $w_B N$ and $w'_B N'$.

#### 4 Gene curation

In this section, we describe how we selected the set of genes that were used for fitting and analysis. The model is designed to fit a single gene, describing the activity of RNA polymerase II through its life cycle. Consequently, for this work we sought a set of isolated genes minimally impacted by overlapping and proximal transcripts. Though there are tens of thousands of genes within the genome, many reside in transcription dense regions– often with overlapping transcripts. Importantly, most genes suffer from one of the following complications: they overlap with the signal from at least one other gene, they have signals from multiple distinct isoforms with differing TSS or PAS coordinates, there exist other types of transcribed loci such as enhancers that are proximal to (or contained within) the gene, or the transcription profiles do not appear to be supported by one or more cell-types. Therefore we curated a subset of genes whose profiles were not obviously impacted by any of these complications, since these complications would confound the interpretation of model fits.

To arrive at this set of genes, we performed several independent computational pre-filters to narrow down the gene list and then inspected these (manually) to confirm that they were free of overlapping transcripts in most of the samples. The first pre-filter consisted of selecting only protein coding genes. Next we took two independent filtering approaches: one based on the distance of a gene to its nearest neighbor, based on the coordinates of their annotations (“annotation approach”) and the other based on a gene’s coverage across all samples relative to the coverage of its nearest neighbor genes (“coverage approach”).

The “annotation approach” consisted of identifying those protein coding genes that had no other annotation within 10kb upstream of the TSS and 30kb downstream from the PAS of its longest isoform. The use of the 10kb/30kb pads was to allow sufficient space for the model to fit run-on transcription signal beyond the bounds of the gene’s annotation and was based on run-on ranges observed previously[4]. This filtering could be done simply from the NCBI hg38.p14 transcript annotations alone (release GCF_000001405.40-RS_2023_03), without the need to consider the coverage files. A single isoform was selected per gene, based on which had the most signal upstream of the $3'$ end of the UTR (a proxy for the PAS). However, this filtering approach fails when there are transcribed loci within the padded window that are unannotated or when the transcription from neighboring loci extend into the 10kb/30kb window. Furthermore, many of of the genes that pass this filter are simply not transcribed in a large subset of the samples and thus cannot be fit. This “annotation approach” filter results in fewer than approximately 1000 genes.

The “coverage approach” consisted of keeping only those genes with an average coverage across all samples greater than 0.1x (over the body of the gene) whose upstream and downstream nearest-neighbor genes had average coverage below 0.1x (to reduce the chance neighboring genes contribute significant signal within the fitting window of the gene being fit). The coverage values were calculated based on the coordinates of the largest isoform for each gene. This filter assured that the gene being considered had sufficient signal across a broad collection of data. However, since the coverage of the neighboring genes was also calculated based on the annotation, this filter failed in cases were the neighboring gene’s transcription signal had significant coverage beyond the bounds of its annotation (e.g. in the case of a prominent dissociation peak significantly downstream from the $3'$ end of its annotation, which may even overlap with the gene being considered). Similar to the “annotation filter,” this filter also could not account for the presence of unannotated, highly transcribed loci proximal to the gene. Genes passing either of these two filtering approaches resulted in about 1400 unique genes.

Using the combined list from computational filtering, each gene was padded by 10kb upstream and 30kb downstream then visually inspected. Manual inspection was necessary because the full extent of transcription is not annotated and neither are most enhancers, thus overlapping transcription could be easily missed by our computational filters. The array of meta samples were loaded into IGV for this visual inspection. Each gene was evaluated based on the following criteria:

1. Does the gene appear to be transcribed in all/most of the meta-samples (see Table 1)?
2. Does the full Loading to Termination profile occur within the padded window?
3. Are there no transcript signals overlapping with the gene of interest that are not a part of the Loading to Termination profile (e.g. upstream/downstream genes that run through the gene being evaluated)?
4. Are there no obvious alternate isoforms, e.g. multiple $5'$ bidirectional peaks or $3'$ peaks, for the gene?
5. Are there no other transcriptional signals in the window, upstream or downstream, that are not associated with the gene of interest (e.g. adjacent genes, eRNA or unannotated lncRNA)?

If the answer to all of these questions was 'Yes' then we would retain the gene with a $5'$end/$3'$end padding length of 10kb/30kb. If the answer to question #5 was 'No' we then determined if we could reduce the size of the $5'$end/$3'$end padding to exclude the additional signal without interfering with the gene's transcriptional profile and, if so, keep it with the adjusted pad lengths. In a few cases, we had to expand the padding beyond the 10kb/30kb to capture the extended dissociation position induced by heat shock. We identified 163 genes from chromosomes 1–6 that were ultimately used for validation and testing (see the gene list in the curated annotation file on the LIET GitHub repo: Dowell-Lab/LIET/resources/chr1-6.liet.ann).

#### 5 Samples

We collected previously published nascent run-on RNA sequencing datasets (PRO-seq) available on the Sequence Read Archive. All samples were processed (QC, trimming, and mapping) during the preparation of the the nascent sequencing database DBNascent[2]. We aimed to get as broad a diversity of human cell/tissue types as possible. Bedgraphs were generated from the processed sample BAM files based on the 3′ end location of each read (typically the end closest to the true polymerase location). The set of samples consisted of both single- and paired-end libraries. Only the 3′ end most read of the paired-end libraries was retained. The following table contains the sample's NCBI accession code (SRR), cell-type, tissue, and experimental condition.

Supplementary Table 1: Cell types, tissues, and accession numbers analyzed in this paper. This analysis was performed across 24 cell types, 12 tissues, and 152 accession numbers. For each celltype, the meta-sample was generated by combining the collection of samples listed in the Accessions column (as described in Methods).

| Cell line | Tissue | Condition | Accessions |
|---|---|---|---|
| A375 | skin epithelial | control | SRR8706163<br>SRR8706164<br>SRR8706165 |
| A549 | lung epithelial | control | SRR12482690<br>SRR12482691 |
| BEAS-2B | lung/bronchus epithelial | control | SRR12482692<br>SRR12482693 |
| CD34 erythroblast | bone marrow | control | SRR11785044<br>SRR11785045<br>SRR11785048<br>SRR11785049 |
| CD4 T cell | bone marrow | control | SRR4012391<br>SRR4012392<br>SRR4012390<br>SRR1810069<br>SRR6780907<br>SRR4012397<br>SRR4012396<br>SRR1810070<br>SRR1810068<br>SRR1810067 |
| ESC | early embryo | control | SRR10669536<br>SRR10669537<br>SRR10669538 |
| HFF | foreskin fibroblast | control | SRR7041686<br>SRR7041689<br>SRR7041690 |
| G401 | kidney epithelial | control | SRR6498830<br>SRR6498831<br>SRR6498832 |
| HAP1 | bone marrow | control | SRR12699335<br>SRR12699334 |

| | | | |
|---|---|---|---|
| HCT116 | colorectal epithelial | control | SRR6290501 |
| | | | SRR6290500 |
| | | | SRR6290499 |
| | | | SRR9304731 |
| | | | SRR9304730 |
| | | | SRR6290518 |
| | | | SRR6290517 |
| | | | SRR6290516 |
| | | | SRR6290515 |
| | | | SRR6290510 |
| | | | SRR6290509 |
| | | | SRR6290508 |
| | | | SRR6290507 |
| | | | SRR6290502 |
| | | | SRR8867628 |
| | | | SRR6290526 |
| | | | SRR6290525 |
| | | | SRR6290523 |
| | | | SRR6290524 |
| | | | SRR8867629 |
| | | | SRR8867637 |
| | | | SRR8867636 |
| HEK293/T | kidney epithelial | control | SRR6026781 |
| | | | SRR6026782 |
| | | | SRR6927839 |
| | | | SRR7988488 |
| | | | SRR7988490 |
| HeLa | cervical epithelial | control | SRR5799635 |
| | | | SRR8478992 |
| | | | SRR8478993 |
| | | | SRR8478996 |
| | | | SRR8478997 |
| K562 | bone marrow | control | SRR12083665 |
| | | | SRR8137173 |
| | | | SRR4454568 |
| | | | SRR4454567 |
| | | | SRR11793826 |
| | | | SRR11793825 |
| | | | SRR12083664 |
| | | | SRR1554311 |
| | | | SRR1554312 |
| | | | SRR5364303 |
| | | | SRR5364304 |
| | | | SRR8669163 |
| | | | SRR8669162 |

| | | | |
|---|---|---|---|
| Kasumi1 | bone marrow | control | SRR3713700<br>SRR5382460<br>SRR5382461<br>SRR12091767<br>SRR12091768<br>SRR5382459<br>SRR5382458<br>SRR5382457<br>SRR5382456<br>SRR3713703<br>SRR3713712<br>SRR3713713 |
| KBM7 | bone marrow | control | SRR10354625<br>SRR10354624<br>SRR10354600<br>SRR10354601<br>SRR10354602<br>SRR10354603<br>SRR10354608<br>SRR10354609<br>SRR10354610<br>SRR10354611 |
| LC2ad | lung epithelial | control | SRR12482694<br>SRR12482695 |
| LCL | blood | control | SRR6727941<br>SRR6727942<br>SRR6727943<br>SRR6727944<br>SRR6727945<br>SRR6727946<br>SRR6727947<br>SRR6727948<br>SRR6727949<br>SRR6727950<br>SRR6727951<br>SRR6727952<br>SRR6727953<br>SRR6727954<br>SRR6727955<br>SRR6727956<br>SRR6727957<br>SRR6727958<br>SRR6727959 |

| | | | |
|---|---|---|---|
| MCF7 | breast epithelial | control | SRR3541129<br>SRR5150536<br>SRR5150537<br>SRR5150538<br>SRR5150539<br>SRR5150540<br>SRR5150543<br>SRR5150546<br>SRR5150549<br>SRR5150552<br>SRR5150555 |
| MEL624 | skin | control | SRR8234171<br>SRR8234172<br>SRR8234173 |
| NUDUL1 | brain | control | SRR8992358<br>SRR8992362 |
| Ramos | blood and bone marrow | control | SRR8544024<br>SRR8544025 |
| SUDHL4 | blood | control | SRR8992356<br>SRR8992360 |
| THP1 | blood | control | SRR11059463<br>SRR11059464 |
| U936 | blood | control | SRR5382467<br>SRR5382468 |
| HeLa | cervical epithelial | w/wo mutant INTS11 (E203Q) treatment | SRR8479004<br>SRR8479005<br>SRR8479006<br>SRR8479007 |
| LCL | blood | w/wo heat-shock treatment (37 and 42°C) | SRR14352131<br>SRR14352132<br>SRR14352135<br>SRR14352136 |

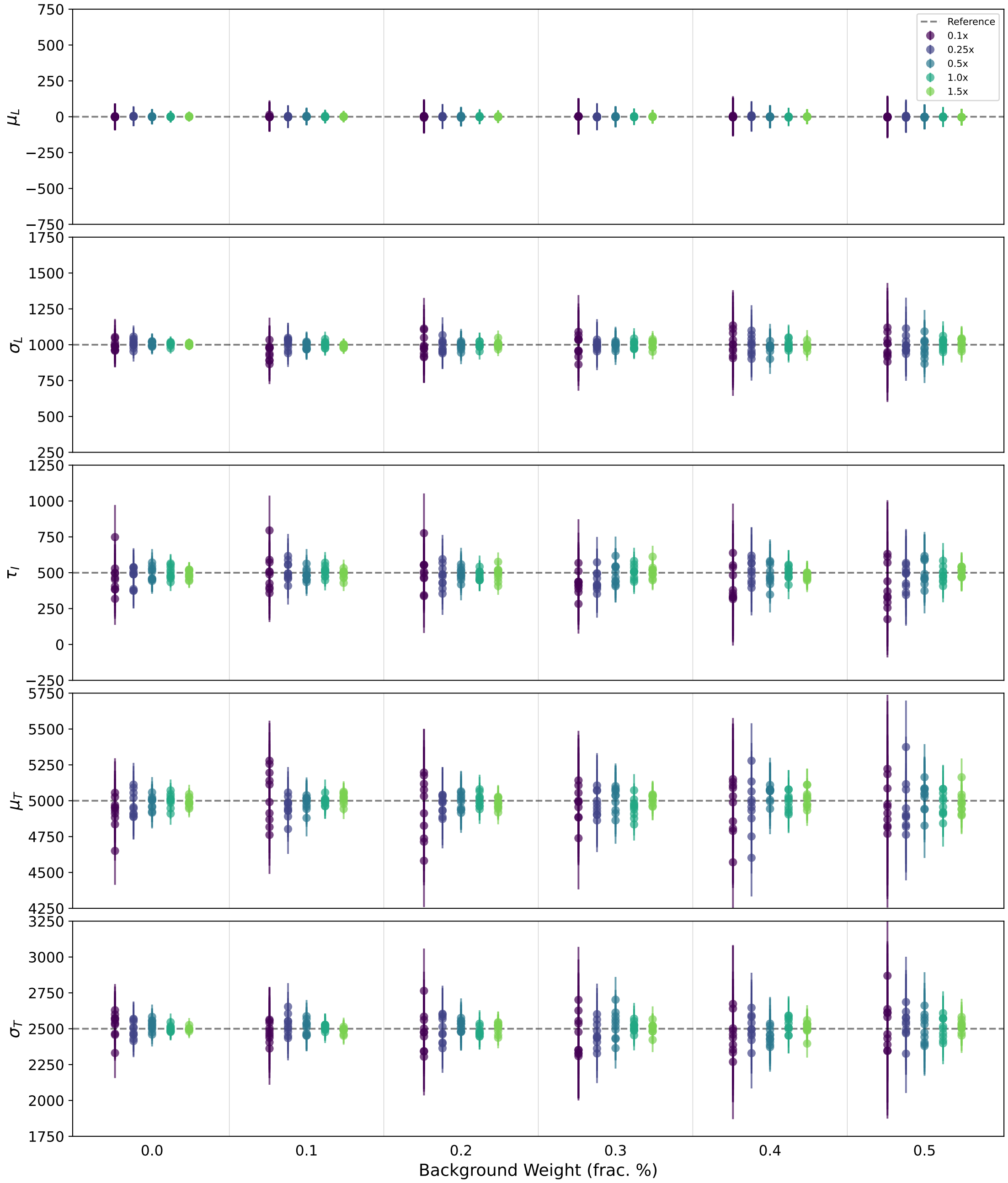


Supplementary Figure 6: Demonstration of the fitting precision/accuracy of the LIET model position and shape parameters as a function of read coverage (legend: purple to green with increasing depth) and varying levels of background (x-axis). We simulated data for a fixed model shape ($\mu_L = 0$, $\sigma_L = 1000$, $\tau_I = 500$, $\mu_T = 5000$, $\sigma_T = 2500$, indicated by the dashed reference lines), representative of a typical gene (length 20kb), with varying levels of total read coverage (shown in the legend: 0.1x $\rightarrow$ 1.5x coverage) and varying proportions of those reads assigned to background (horizontal axis: 0% $\rightarrow$ 50%). For each combination of coverage and background, ten *i.i.d.* simulations were generated and fit with the LIET model. Each point represent the best-fit estimate for the specified parameter (y-axes) for a single simulation fit and the error bar on the point represents $3\sigma_{post}$ for that fit (see discussion in Supp. Sec. 2).

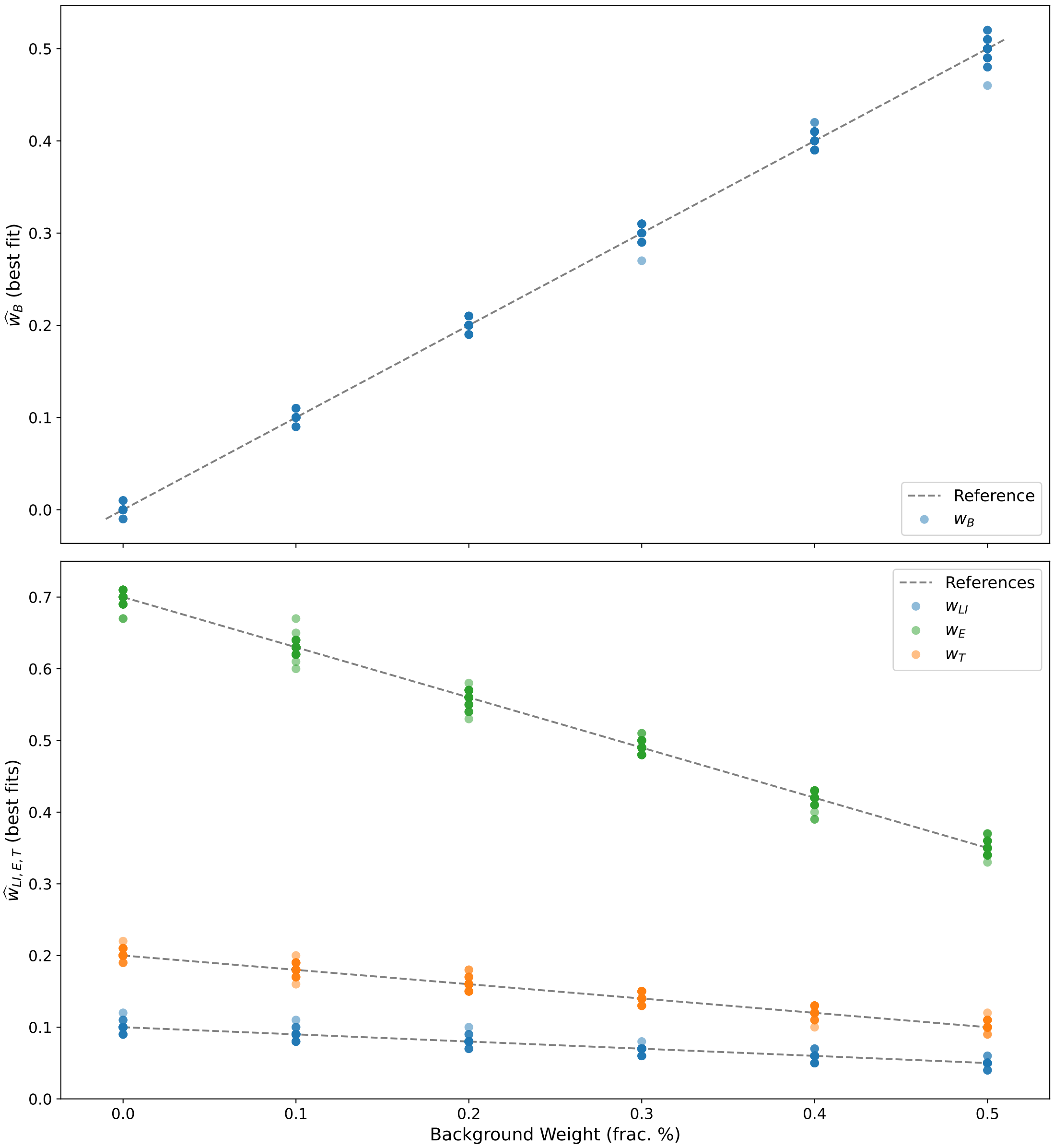


Supplementary Figure 7: Demonstration of the fitting precision/accuracy of the LIET model component weights (legend) as a function of read coverage and varying levels of background (x-axis). These are the best-fit estimates for the model weights for the same simulations as those in Supp. Fig. S.6 as a function of proportion of background reads (horizontal axis). **Top:** The best-fit values for the background weight component ($w_B$, blue points) compared to the simulation reference (dashed line). The variation $w_B$ is extremely low such that most of the points overlap one another (there are 50 simulations for each background value: 10 *i.i.d.* simulations for each of the 5 different total coverage values). In other words, the precision of the fit is robust to variation in coverage. **Bottom:** The best-fit values for the LI, E, and T components of the model ($w_{LI}$, $w_E$, $w_T$, respectively) as a function of background weight. Just as with the $w_B$ estimates, these reproduce the reference values with very high precision. Notes: Rounding precision for the component weights was set to 0.01. The bottom point colors correspond to the model components in Fig. 1 of the main text.

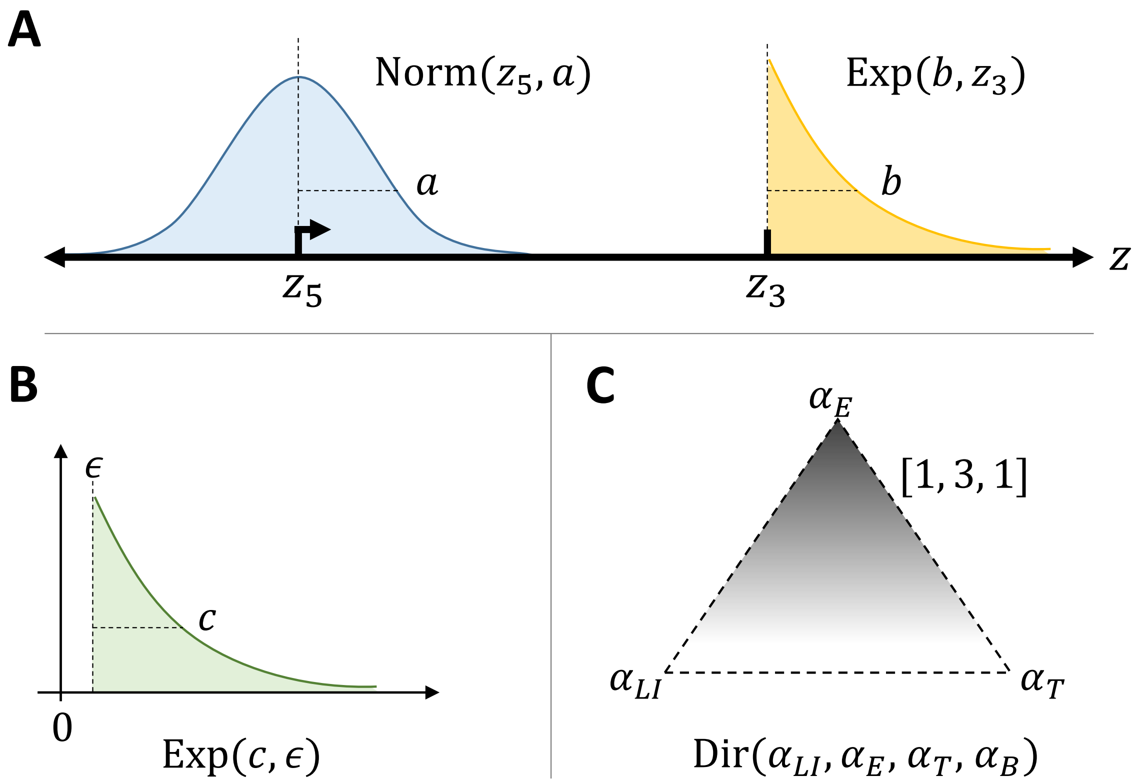


Supplementary Figure 8: The types of prior distributions ($p(\boldsymbol{\Theta})$) used for the model parameters. (A) Those used for the location parameters $\mu_L, \mu'_L$ are normal distributions (blue) with width $a$, which are centered on the $5'$ reference point $z_5$ (typically equal to the gene's TSS). The prior distribution for the location parameter $\mu_T$ is an exponential (yellow) with characteristic length $b$ and is located at the $3'$ reference point $z_3$ (typically the end of the gene's $3'$-most exon). In other words, the value of $z_3$ sets the upper bound for the optimization of $\mu_T$. Genomic coordinate is indicated by $z$. (B) Exponential distributions are also used for the shape parameters $\sigma_L, \tau_I, \sigma_T, \sigma'_L, \tau'_I$. In principle, the value of the characteristic exponential length $c$ is different for each parameter (see Methods in the main text for the values used for each) and the location value $\epsilon$ is equal to zero (but may be set to a small, non-zero value to set a lower bound on the value of each shape parameter). (C) The prior distribution for the model component weight parameters are a 4-dimensional Dirichlet distribution (three model and one background component). Here we show an example for the three model components with unequal concentration hyper-parameters ($[\alpha_{LI}, \alpha_E, \alpha_T] = [1, 3, 1]$), demonstrating a prior where *Elongation* would be more likely. For all fitting we used $[\alpha_{LI}, \alpha_E, \alpha_T, \alpha_B] = [1, 1, 1, 1]$.

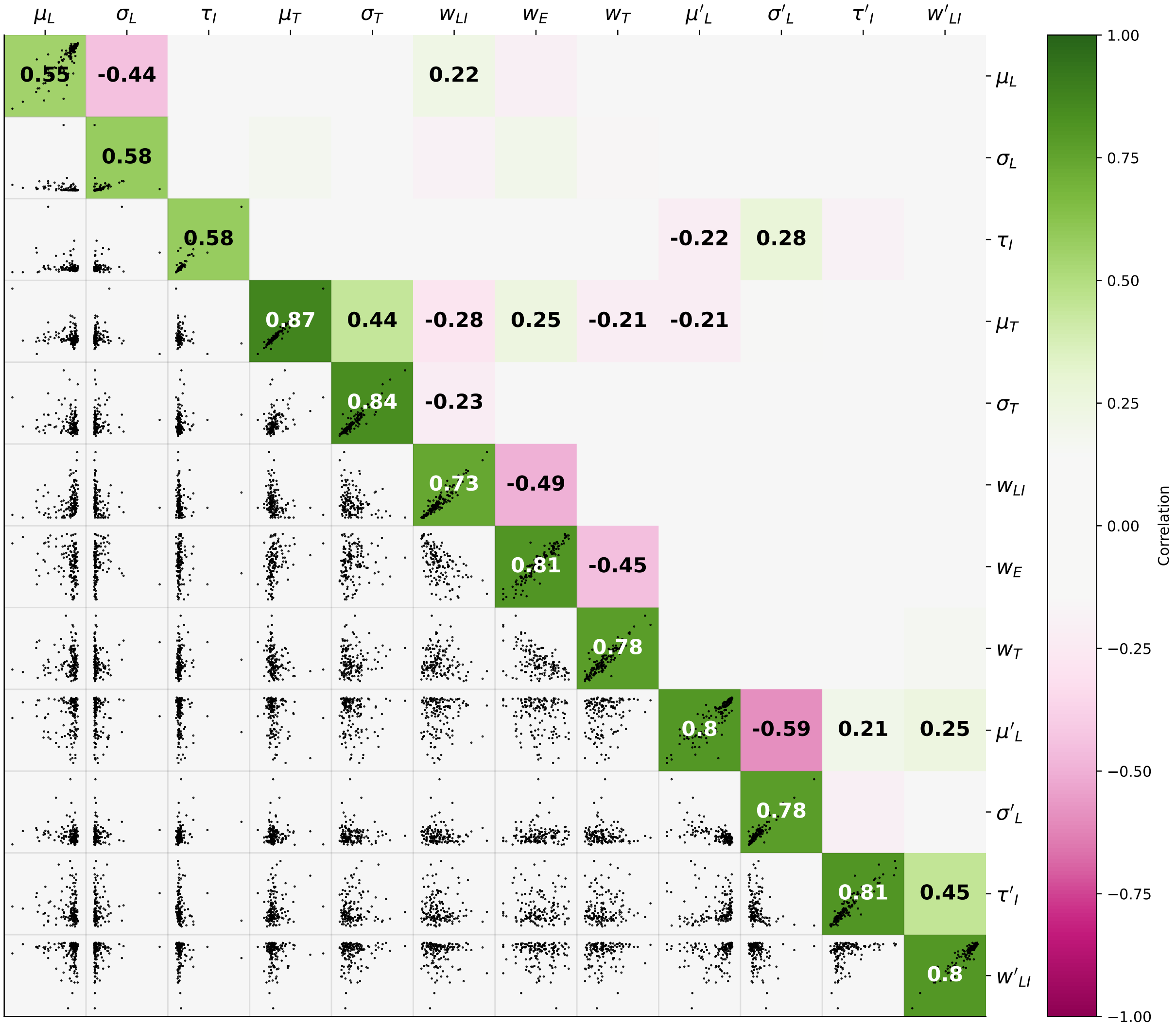


Supplementary Figure 9: Upper triangle is a reproduction of the cross-parameter correlation analysis in Fig. 2Cwhich was run on ten control replicates of HCT116 data from three papers. Here the lower triangle depicts the cross-parameter comparison scatter plots (each point a gene from our list) of the two replicates whose correlation is the closest to the median correlation value. The numeric values are the median correlation coefficients across all pairwise comparisons between samples. Only those median correlation coefficients with magnitude $> 0.2$ are labeled.

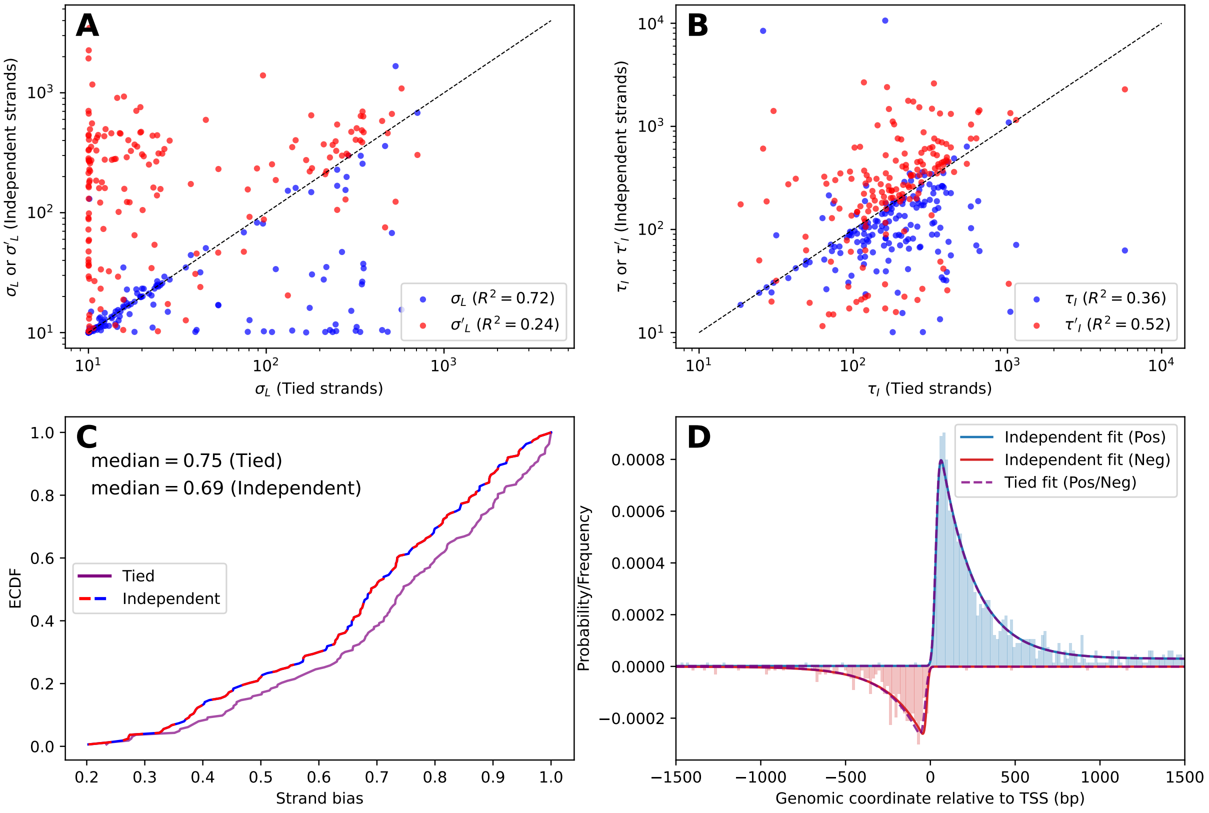


Supplementary Figure 10: Supplement to Fig. 3, in the main text. (A) Scatter plot comparing $\sigma_L$ (blue) and $\sigma'_L$ (red) fit results under the independent fitting scenario to that of the tied strand scenario. Note that $\sigma'_L$ is systematically larger than $\sigma_L$, when fit independently. Dashed line is one-to-one line. The $R^2$ values indicate $\sigma_L$ predominates under the tied scenario. (B) Scatter plots comparing $\tau_I$ (blue) and $\tau'_I$ fit results under independent scenario to that of the tied scenario. (C) Empirical cumulative distribution function of the strand bias (Eq. 14) under the independent (blue-red-dashed) and tied strand (purple) fitting scenarios. The strand bias is computed using the shape parameters from A and B (and weights) under the two scenarios. (D) Schematic of generated data from the median 5′ parameter values from the tied scenario, fit by the tied (purple) or independent (blue/red) scenario. Note that both scenarios fit well to the generated data, in contrast to the converse simulation, which was fit better by the independent scenario.

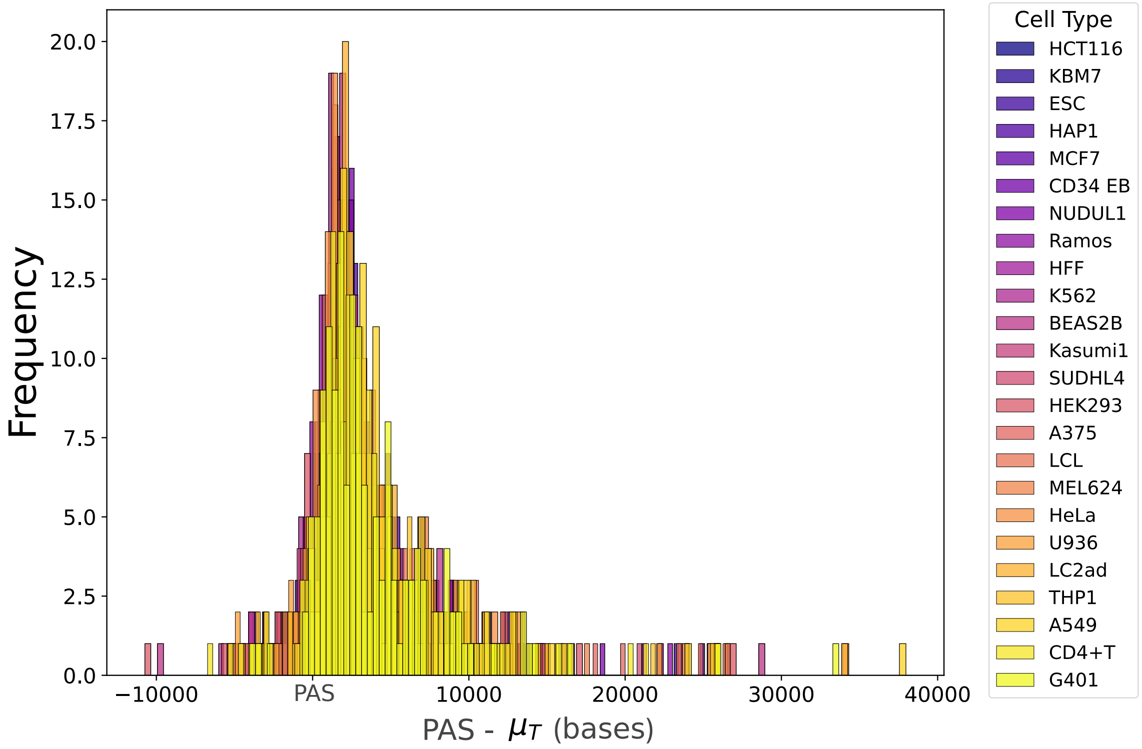


Supplementary Figure 11: Histogram representation of the PAS to $\mu_T$ distances for the genes in our gene list in all control-condition cell-types. This data is equivalent to that presented in the heat map in Fig. 4A. The distributions are statistically equivalent between all cell-types.

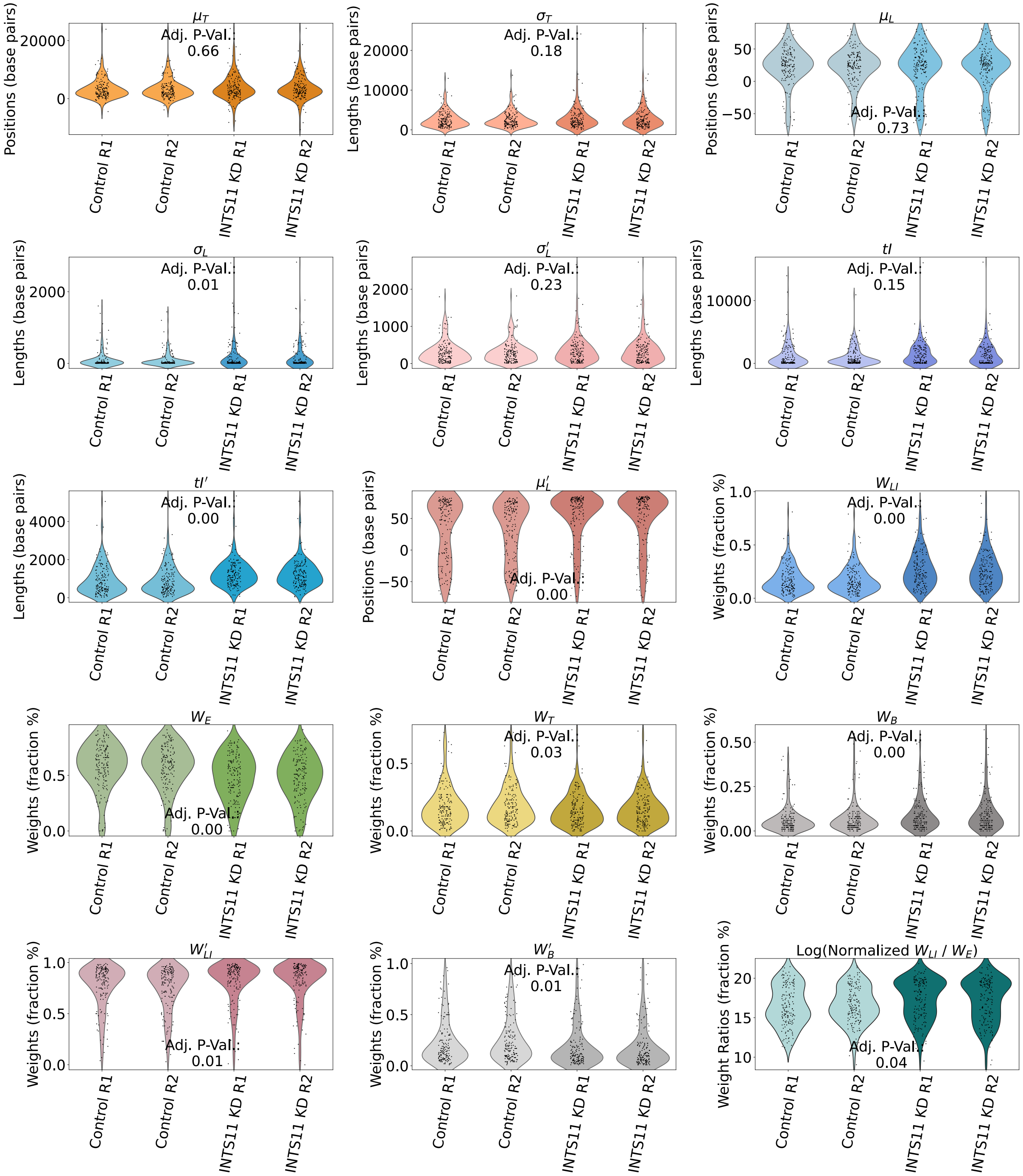


Supplementary Figure 12: Supplemental violin plots for all position, shape, and weight parameters for the INTS11 knock-down experiment presented in Fig. 5D–F, in the main text. The p-adj values indicate the one-way pairwise ANOVA comparison between control and INTS11 knock-down. In this INST11 knock-down experiment[5], Beckendorff et al. generated HeLa cells with stable doxycycline-inducible (dox) short hairpin RNA against the INTS11 subunit of the integrator catalytic module. Both control and experimental samples, shown in Fig. 5 of the main text, from this experiment were treated with the exogenous E203Q mutant. However, only INTS11 knock-down labeled samples were treated with doxycycline, enabling the shRNA INTS11 knock-down.

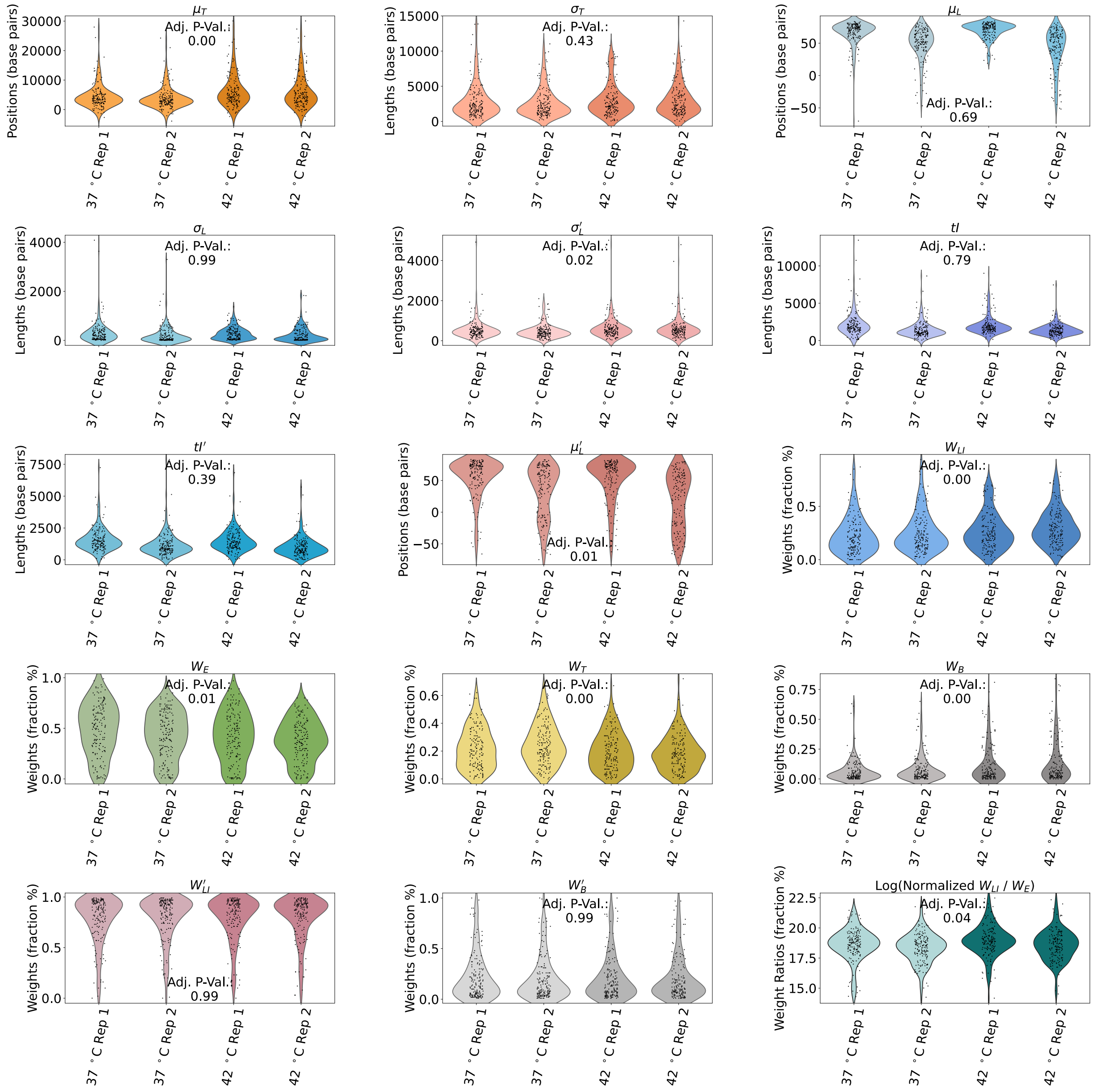


Supplementary Figure 13: Supplemental violin plots for all position $(\mu_L, \mu_T, \mu'_L)$, shape $(\sigma_L, \tau_I, \sigma_T, \sigma'_L, \tau'_I)$, and weight parameters for the heat shock experiment presented in Fig. 5A–C, in the main text. The p-adj values indicate the one-way pairwise ANOVA comparison between control and heat shock condition.
